# BRAF^V600E^-Driven Lung Tumorigenesis Requires Ligand-Mediated Activation of ERBB Receptor Signaling

**DOI:** 10.1101/2025.05.04.652129

**Authors:** Melanie Angelina Dacheux, Meng-Jung Wu, Michael T. Scherzer, Monique Nillson, Brandon Murphy, Sophia Schuman, Zhenlin Ju, Trever Bivona, Piro Lito, Matthew Gumbleton, Sonam Puri, Wallace Akerley, Jing Wang, Kaiwen Wang, John Heymach, Conan G. Kinsey, Marcelo Negrao, Martin McMahon, Aria Vaishnavi

## Abstract

Secretion of ligands of the human epidermal growth factor (EGFR) family of receptors or erythroblastic leukemia viral oncogene family (ERBB1-4) is a feature common to many cancer cells. However, our understanding of the role of autocrine ligands in the aberrant behavior of cancer remains incomplete. Here we demonstrate that, in numerous preclinical models of lung tumorigenesis, BRAF^V600E^ signaling promotes expression of ligands including *HB-EGF*, *TGFα*, *Epi-* and *Amphiregulin*. Moreover, using both genetic or pharmacological approaches, we demonstrate that ligand-mediated activation of EGFR signaling in the tumor cell is required to sustain both early-stage BRAF^V600E^-driven lung tumorigenesis and supports late-stage BRAF^V600E^-driven lung cancer maintenance. Unbiased Reverse Phase Protein Analyses (RPPA) analyses, paired with targeted validation, reveals ERBB signaling serves to sustain signaling through the ERK1/2 MAP kinase pathway, through effects on ARAF and CRAF, and on the parallel JUN kinase (JNK) pathway. Furthermore, EGFR is activated in a cohort of *BRAF*-mutated lung cancer patients both pre- and post-treatment. Finally, we noted significant improvement in the depth and durability of therapeutic responses in preclinical models of BRAF^V600E^-driven lung cancer by combined inhibition of both BRAF^V600E^ signaling plus pan-ERBB signaling. Collectively, this work provides evidence for an important role for ERBB family signaling in the genesis and maintenance of BRAF^V600E^-driven lung cancers, and the potential for future therapeutic improvement by rational combination targeting of these pathways.

**SIGNIFICANCE:** *BRAF^T1799A^* serves as a predictive biomarker for FDA-approved targeted inhibition of BRAF^V600E^ oncoprotein kinase signaling in non-small cell lung cancer (NSCLC). However the occurrence of primary or acquired drug resistance limit the depth and durability of patient responses. Studies described here provide a mechanistic rationale for clinical testing of first-line BRAF^V600E^ inhibition combined with pan-ERBB inhibition to improve the depth and durability of initial patient responses, and delay the emergence of lethal drug resistant disease.

## INTRODUCTION

Non-small cell lung cancer (NSCLC) is the leading cause of cancer-related deaths globally, with adenocarcinoma being the most common sub-type^1^. Mutationally-activated KRAS or BRAF are actionable oncoprotein drivers of lung adenocarcinoma, acting to elevate flux through the RAS-regulated RAF>MEK>ERK mitogen activated protein kinase (MAPK) signaling pathway, which is one of the most commonly dysregulated pathways in cancer. Hot-spot *BRAF* mutations occur in ∼8% of NSCLC (representing ∼ 20,000 patients in the U.S. annually). Importantly, it has been estimated that the same number of patients die annually of BRAF^V600E^ melanoma and BRAF^V600E^ lung cancer, reaffirming the relevance of this cancer subset ^2, 3^. Although pathway-targeted inhibitors of BRAF oncoprotein kinase signaling (dabrafenib plus trametinib or encorafenib plus binimetinib) are FDA approved for NSCLC, problems such as intrinsic or acquired drug resistance limit the depth and durability of responses of NSCLC patients to these agents ^4, 5^. Previous work has also revealed that inhibition of BRAF^V600E^ alone is not a sufficient approach in BRAF^V600E^+ colorectal cancer due to the activation of EGFR signaling in these tumors ^6, 7^. As such, a critical gap remains in understanding the tissue-specific signaling mechanisms that confound BRAF^V600E^ inhibitors, particularly in the lung microenvironment. Improving our knowledge of potential tumor progression drug targets in the context of BRAF-driven lung cancer subsets will likely result in an improvement of therapeutic rationale for these patient populations. In this study, we focus primarily on the BRAF^V600E^ subset, but also reveal the commonality of these mechanisms amongst additional MAPK pathway oncogenic drivers of lung cancer.

EGFR was first proposed as a potential lung cancer target over 40 years ago ^8^. While early efforts were focused on targeting normal EGFR signaling in unselected patients or all-comers ^9–11^, the subsequent recognition of mutational activation of *EGFR* (e.g. deletions in exon 19 or a point mutation in exon 21 encoding EGFR^L858R^) provided both a rational and a predictive biomarker for the effectiveness of pharmacological inhibitors of EGFR’s tyrosine kinase activity (e.g. gefitinib or erlotinib) inhibitors in a subset of NSCLC patients ^12, 13–15^. Over the ensuing years, the use of more potent and selective EGFR inhibitors (e.g. osimertinib) in earlier stages of the disease has transformed the care of *EGFR* mutated NSCLC patients from the days of platinum-based chemotherapy ^15^.

The RAS-activated RAF>MEK>ERK signaling pathway is one of the most commonly dysregulated signal transduction pathways in cancer. This pathway has profound effects on numerous biochemical and biological processes that are mission-critical to cancer cell survival ^16, 17^. Many such effects are mediated by transcriptional regulation of mRNA expression, often through AP1 or ETS family transcription factors ^18, 19^. Among the most prominent transcriptional targets of RAS or RAF signaling are members of the family of ligands of EGFR and its close family of ERBB receptors such as HB-EGF, TGFα and amphiregulin ^20, 21^. This data suggests that constitutive activation of RAS>RAF>MEK>ERK signaling can activate the 4 ERBB family members in an auto-, juxta- or paracrine manner. Previous work has also revealed that oncogenic KRAS^G12D^ relies on trophic signaling from the ERBB family in lung cancer, and that pan-ERBB inhibition may be an important target in KRAS-driven lung cancer ^22, 23^. However, the full molecular and functional consequences of ERBB family signaling have not been elucidated clearly in the context of BRAF^V600E^ in lung cancer. Here, we set out to more deeply elucidate the molecular mechanisms that regulate signaling cross talk between BRAF^V600E^ oncoprotein kinase signaling and the ERBB family of receptor tyrosine kinases, and conclude that BRAF^V600E^-driven lung tumors rely on ERBB signaling for the initiation of early-stage tumor growth and the maintenance of late-stage lung cancers.

## RESULTS

### BRAF^V600E^ regulates transcription of ERBB ligands *in vivo* in AT2 cells

We began our study by investigating the transcriptional effects of BRAF^V600E^ signaling *in vivo* through targeted BRAF^V600E^ inhibition with the FDA-approved therapeutic combination of dabrafenib and trametinib against various mouse models of BRAF^V600E^. Here, the following cohorts of mice were initiated: *Braf^V600E^*+ or *Braf<u>^C^</u>^re-<u>A</u>ctivatedD^ ^td<u>T</u>omato^ ^(CAT)^* genetically engineered mouse model studies: *Braf^CAT^* alone (*B*), *B* + *Trp53^floxed^* (*BP*), *B* + *CDKN2A or Ink4a/Arf^FloxedD^* or (*BC*), *B* + *Trp53^R172H^* or (*BP172*), *B* + *Trp53^R245W^* or (*BP245*), or *B* + *Pik3CA^1057^* or (*BP1057*). Lung tumors were initiated by intranasal instillation of 2.5 x 10^7^ pfu of adenovirus encoding CRE recombinase (Ad5.SPC-CRE) for 8 weeks, prior to once daily treatment with either vehicle or 75 mg/kg <u>D</u>abrafenib with 1 mg/kg <u>T</u>rametinib (D+T). Following 1-2 weeks of treatment, mice were euthanized and lungs were processed for single cell sequencing analyses on the 10x Genomics platform. Here, we leveraged a tdTomato reporter that runs off the endogenous *Braf* promoter in the *Braf^CAT^* mouse paired with flow cytometry following lung digestion to enrich our isolation of tumor cells for these analyses ^24, 25^. Transcriptomic analyses revealed a clear trend that all 7 EGFR ligands were present or harbored high expression levels in vehicle treated lungs, but decreased following treatment with D+T (**Figure 1A**). These data also demonstrated that the expression of EGFR ligands, particularly *Areg*, *Hbegf*, *Ereg*, and *TGFa*, are controlled by BRAF^V600E^ in an autocrine manner. This conclusion is based on the induction of these lung tumors by the *Surfactant Protein C* (SPC) promoter, a marker of alveolar type 2 (AT2) pneumocytes (the putative cell of origin for lung adenocarcinoma) paired with the isolation of tdTomato+ AT2 cells for these *in vivo* analyses. Further biochemical interrogation through immunoprecipitation and immunoblot analyses revealed BRAF^V600E^ signaling also regulates expression of the EGFR ligand HB-EGF in human lung cancer cells, as well (Supplementary Figure 1A).

**Figure 1.**
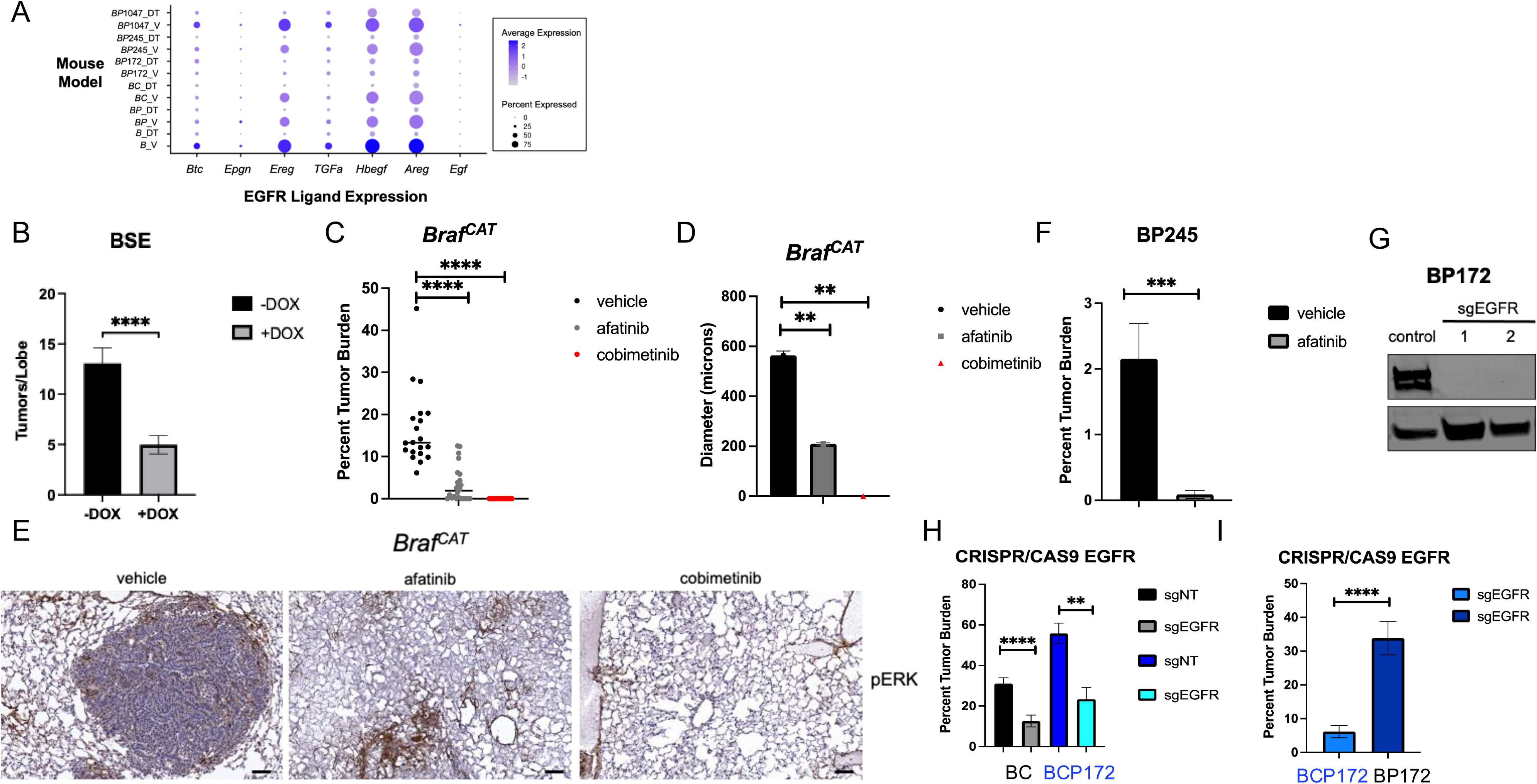
Pharmacological and Genetic Blockade of ERBB Signaling Impairs the Initiation and Progression of BRAF^V600E^-Driven Tumors (GEMMs). A.) Transcript levels of 7 EGFR ligands extracted from single cell sequencing analyses from vehicle (V) or 75 mg/kg dabrafenib + 1 mg/kg trametinib (DT) treatment of the following alleles *Braf^V600E^*+ or *Braf<u>^C^</u>^re-<u>A</u>ctivatedD^ ^td<u>T</u>omato^ ^(CAT)^* genetically engineered mouse model studies: *Braf^V600E^* alone (*B*), *B* + *Trp53^floxed^* (*BP*), *B* + *CDKN2A or Ink4a/Arf^FloxedD^* or (*BC*), *B* + *Trp53^R172H^*or (*BP172*), *B* + *Trp53^R245W^* or (*BP245*), or *B*+ *Pik3CA^1057^* or (*BP1057*). Each allele was initiated with 2.5 x 10^7 PFU Ad5mSPC-CRE 8 weeks prior to starting pharmacological intervention. Mice were dosed once daily for up to two weeks by oral gavage with V or DT prior to euthanasia for single cell analyses of lung tissue. B.) Quantification of tumors/lobe in *Braf^CAT^* + *SPC*-rtTA-*EGFR-tr* (BSE mice) either + or – doxycycline chow after 10- or 12-weeks post-initiation. Error bars represent SEM. *** = p < 0.001 by an unpaired Welch’s T-Test. N=15 mice (-DOX) or 14 mice (+DOX), respectively, in 2 unique experiments. C.) Quantification of tumor burden in *Braf^CAT^* or B mice treated immediately post initiation once daily via oral gavage of vehicle control, 5 mg/kg cobimetinib (positive control), or 15 mg/kg afatinib for 8 weeks. Error bars represent SEM. **** = p < 0.0001 by a one-way ANOVA. N=7 mice vehicle, 7 mice afatinib, 6 mice cobimetinib. D.) Quantification of tumor diameter of same mice previously described in (C). Error bars represent SEM. ** = p < 0.01 by a one-way ANOVA. E.) Immunohistochemistry analyses of phosphorylated ERK (pT202, Y204) in *Braf^CAT^* formalin fixed paraffin embedded (FFPE) mouse lung tissue described in (C). F). Quantification of tumor burden in *Braf^CAT^*; *Trp53^R245W^* or BP245 mouse lungs at 8 weeks post initiation following once daily oral gavage with vehicle control or afatinib. Error bars represent SEM. ** = p < 0.01 by an unpaired Welch’s T--Test. N=16 mice (9 vehicle or 7 afatinib per group). G.) Immunoblot analyses of EGFR expression following stable expression of vector control, or 2 unique sgRNA against mouse EGFR in a BP172 cell line. H.) Quantification of lung tumor burden in *BC* (*Braf^CAT^*^;^ *H11^LSL-CAS9^*) or *BCP172* (*Braf^CAT^*^;^ *H11^LSL-CAS9^ ;Trp53^R172H^*) mice initiated with lentiviruses encoding either non-targeting control sgRNA or sgEGFR and Cre Recombinase. Error bars represent SEM. **** = p < 0.0001 or ** = p < 0.01 by a one-way ANOVA. N=20 (5 per group). I.) Quantification of lung tumor burden in BCP172 (*Braf^CAT^*^;^ *H11^LSL-CAS9^ ;Trp53^R172H^*) or BP172 (*Braf^CAT^*^;^ *Trp53^R172H^*) mice initiated with lentivirus encoding sgEGFR after 8 weeks. Error bars represent SEM. **** = p < 0.0001 by an unpaired Welch’s T-Test. N=25 mice.

### Pan-ERBB Signaling Is Necessary for BRAF^V600E^ Lung Cancer Initiation *In Vivo*

Next, we set out to determine the functional relevance of these EGFR ligands in BRAF^V600E-^ driven lung tumorigenesis. To test this, we deployed a genetic EGFR ligand trap using a combination of two mouse transgenic alleles: *SPC*-*rtTA* driving expression of *EGFR_trunc_*-*tetO* ^26, 27^. In this model, doxycline (DOX) induces expression of the reverse tetracycline transactivator (rtTa) from the SPC locus, which activates a truncated, C-terminal intracellular deletion of EGFR specifically in AT2 pneumocytes (Supplementary Figure 1B, C). Here EGFR is able to bind or trap the ligand but is unable to signal or asymmetrically dimerize, operating in a dominant negative manner as well as a ligand trap. The EGFR ligand trap was combined with *Braf^CAT^* mice to assess the role of EGFR family ligands in promoting lung cancer initiation (Supplementary Figure 1D). A cohort of BES mice (*<u>B</u>raf^CAT^*; *<u>E</u>GFR_trunc_*-*tetO*, *<u>S</u>PC*-*rtTA*;) were initiated with CRE Recombinase expressing virus, immediately placed on DOX chow, and the lungs were assessed at different time points post euthanasia. The activation of the EGFR ligand trap led to a striking reduction in lung tumor burden, revealing a critical, autocrine role for EGFR family ligands in promoting BRAF^V600E^ lung cancer initiation in AT2 cells (**Figure 1B**, *p*<0.0001). To test the relevance of EGFR ligands in other lung cancer driver models, we expanded this tool and tested our ligand trap in an established mouse model of EML4-ALK-driven lung cancers (Supplementary Figure 2A) ^28^. Similarly and consistent with BRAF^V600E^, we saw a robust decrease in tumor burden when the ligand trap was activated, reaffirming the importance of EGFR ligands in the initiation of EML4-ALK-driven tumors as well (Supplementary Figure 2B&C).

Based on these striking genetic results, we next hypothesized that EGFR kinase activity, which drives its signaling functions, is necessary for its role in supporting BRAF^V600E^-driven lung cancer initiation. To test this hypothesis, we switched to using pharmacological inhibition of pan-ERBB signaling with the small molecule tyrosine kinase inhibitor (TKI) afatinib during BRAF^V600E^- driven lung cancer initiation. Importantly, afatinib is a potent wild-type EGFR inhibitor, but can also prevent promiscuous heterodimerization amongst the other ERBB family members. A cohort of *Braf^CAT^* mice were initiated with SPC-CRE and immediately dosed once daily with vehicle, afatinib, or the MEK1/2 inhibitor cobimetinib (a positive control for blocking BRAF^V600E^ signaling) for 8 weeks ^29^. Consistent with the results from the genetic ligand trap, pharmacological blockade of pan-ERBB signaling with afatinib significantly reduced lung tumor burden, and tumor diameter (**Figure 1C&D;** *p*<0.0001 and *p*<0.001). We next reasoned that perhaps activation of a lung cancer hot spot, gain-of-function P53 mutation, *Trp53^R245W^*, may be sufficient to overcome the need for EGFR signaling at lung cancer initiation. However, *BP245* mice dosed with afatinib for 8 weeks still showed a similar a significant reduction in tumor burden by 8 weeks (**Figure 1E&F**; *p* <0.001). To understand the full contributions of mouse EGFR, including functions that could be kinase-independent, we used CRISPR/Cas9 genome editing to silence its protein expression. This loss-of-function (LOF) approach was key, because it allowed us to elucidate any potential roles of EGFR through a scaffolding or kinase-independent based mechanism mediated by the presence of EGFR protein. As such, sgRNAs against EGFR were deployed *in vitro* to test if we could establish EGFR null cells in BRAF^V600E^-driven lung cancers (**Figure 1G**). Subsequently, we observed a clear loss of EGFR protein expression through immunoblot analyses (**Figure 1G**). To test the role of sgEGFR *in vivo*, lentiviruses that encode either a non-targeting control sgRNA (sgNT) or sgEGFR alongside CRE recombinase were administered intratracheally to initiate *Braf*^V600E^; *H11*^LSL-CAS9^ mice with or without *Trp3*^R172H^ (*BC*, *BCP*, *BP* mice). We observed a significant loss in lung tumor burden in mice that received sgEGFR compared to sgNT virus in *BC*, *BCP*172, or *BP*172 mice (**Figure 1H&I**; *p* <0.0001, *p* < 0.01, and *p* < 0.001). Collectively, this data reveals an essential regulatory role for EGFR in supporting BRAF^V600E-^driven lung cancer early on at lung tumor initiation.

To determine whether the functional importance of EGFR signaling at lung tumor initiation was permanent or reversible, we deployed an inducible, conditional RNA interference (RNAi) strategy with DOX regulated short hairpin RNAs (shRNAs) that genetically target mouse EGFR^30^. Here, lentivirus encoding either a control inducible, but reversible shRNA (Renilla or REN) or mouse EGFR was used to create stable BRAF^V600E^ ^or^ EML4-ALK expressing lung cancer cell lines (**Figure 2A&C**, and Supplementary Figure 3A). 24 and 48 hours of EGFR knockdown resulted in a clear reduction in pERK protein levels, an established readout for RAS-regulated RAF>MEK>ERK signaling pathway activity (**Figure 2A&C.**) We hypothesized that EGFR signaling may be important at different stages of tumor growth, and tested this by subcutaneously implanting shREN or shEGFR expressing BRAF^V600E^ (BP245) or EML4-ALK (EAV; EA1M) cells into the flanks of immunocompromised mice in a xenograft model. To begin, shREN or shEGFR mice were randomized on or off DOX chow to test the role of EGFR at initiation of tumor formation in each of these cell line xenograft model systems. While shREN had no consequences on tumor formation, the mice on DOX that were implanted with expressed shEGFR revealed significantly lower tumor volume compared to the paired off DOX controls (**Figure 2B&D**, Supplementary Figure 3B; *p* < 0.0001). Next, to determine whether the effect could be rescued or reversed, we swapped the DOX doses for the two shEGFR expressing arms at day 12 or day 10 until day 21 or day 17, respectively for BP245 and EAV. There was a rapid shrinkage of established tumors that were off DOX, or rescued tumor growth in the previously impaired tumors that were on DOX (*p* < 0.0001), This data reveals that in these models modulation of EGFR expression can reversibly contribute to tumor maintenance. To push these dynamic effects further, we did a final switch to pharmacologic treatment with afatinib after tumors stopped responding to genetic manipulation. Tumors responded one more time to afatinib briefly before finally giving in to acquired drug resistance and tumor progression (**Figure 2B&D**, Supplementary Figure 3B). Furthermore, immunohistochemical analyses of FFPE tumors analyzed post necropsy revealed substantial reduction in pERK staining following afatinib treatment in the EA1M tumor model compared to control (Supplementary Figure 3C). This data demonstrates the spatio-temporal importance of EGFR signaling at tumor initiation and maintenance in multiple models of BRAF^V600E^ and EML4-ALK-driven lung cancer mouse models, and that these effects are readily reversible to genetic or pharmacological exploitation.

**Figure 2.**
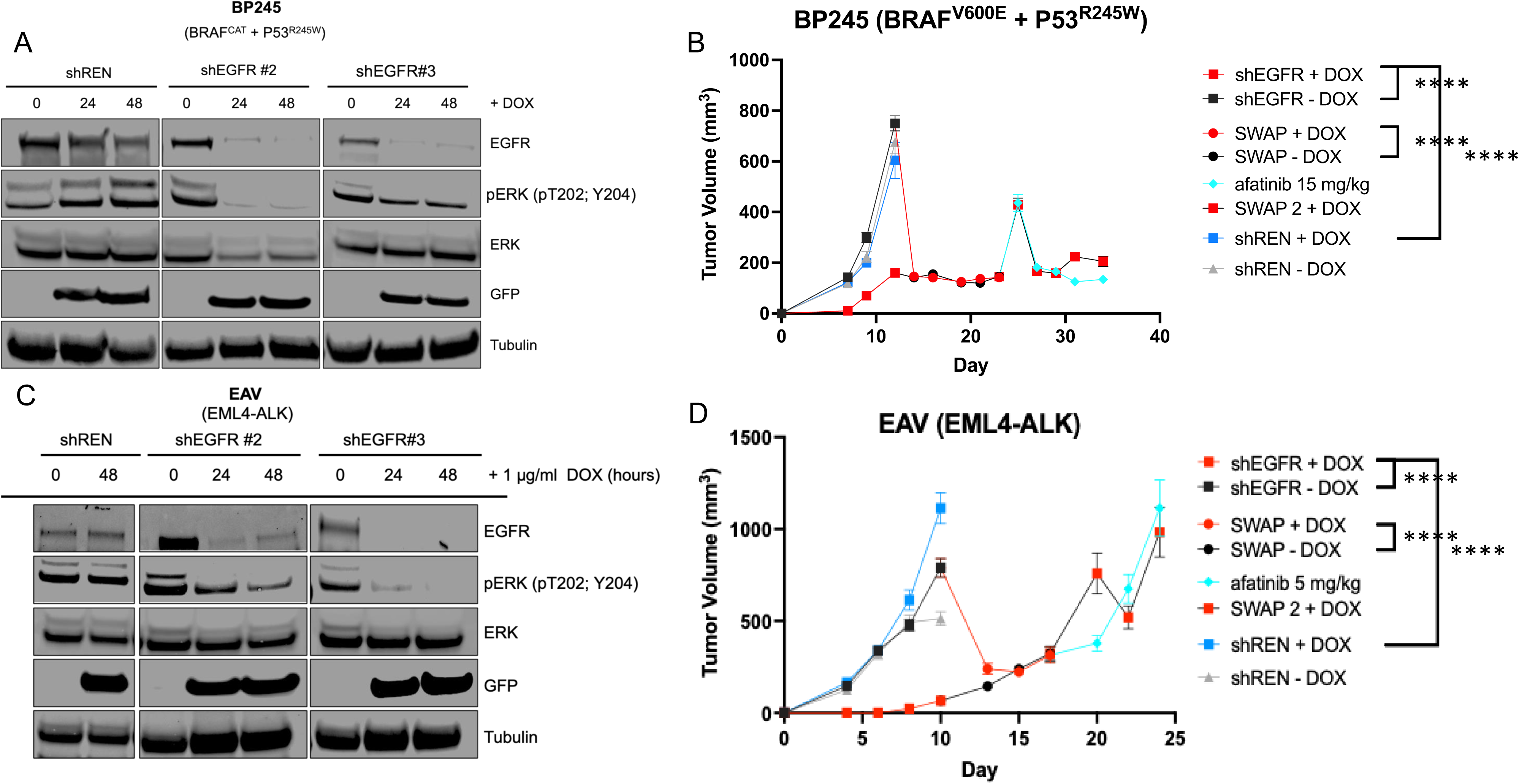
Lung Cancer Xenografts Demonstrate Exquisite Sensitivity to the Anti-Tumor Activity of DOX-Inducible shRNAs Against Mouse EGFR at Different Stages of Tumorigenesis. Immunoblot analyses assessing protein expression with the indicated antibodies in BP245 cell lysates that stably express lentiviruses that encode short hairpin RNAs (shRNAs) against Renila (control) or EGFR treated with 1μg/ml doxycline for 0, 24, or 48 hours. A.) Quantification of tumor volume changes over time in BP245 cells stably expressing control (shREN) or shEGFR expression plasmids subctutaneously implanted into NOD-SCID mice as a xenograft. Mice were on or off doxycycline chow at the indicated timepoints and swapped at the indicated points. For shEGFR expressing xenografts red represents on DOX and black represents off DOX chow. Mice were also dosed with 15 mg/kg afatinib at the indicated times (colored in aqua). Error bars indicate the SEM. **** = p < 0.0001 by a paired T-Test. N= 5-10 mice per group. B.) EAV (EML4-ALK) cell lysates that stably express lentiviruses that encode short hairpin RNAs (shRNAs) against Renila (control) or EGFR treated with 1μg/ml doxycline for 0, 24, or 48 hours. C.) Quantification of tumor volume changes over time of EAV cells stably expressing control (shREN) or shEGFR expression plasmids subcutaneously implanted into NOD-SCID mice. Mice were on or off doxycycline chow at the indicated timepoints and swapped at the indicated points. For shEGFR expressing xenografts red represents on DOX and black represents off DOX chow. Mice were also dosed with 15 mg/kg afatinib at the indicated times (colored in aqua). Error bars indicate the SEM. **** = p < 0.0001 by a paired T-Test. N=5 or 10 mice per group (5 for shREN, 10 for shEGFR – and + DOX).

### Pan-ERBB signaling is an actionable target that contributes to tumor maintenance in BRAF^V600E+^ human xenograft models

The GEMM data was compelling, but we wanted to test whether ERBB signaling was contributing to tumor maintenance in a cohort of human-derived models of patient-derived xenografts (PDX) established from BRAF^V600E^+ human lung cancer tumors (**Table 1**): CTG0167 (**Figure 3A**), NCI349418 (**Figure 3B**), HCIBL1 (**Figure 3C**) , Lito7S (**Figure 3D),** and Lito1S (**Figure 3E&F**). To that end, we initiated all 5 PDX models via subcutaneous implantation into immunocompromised mice, and randomized them onto 3 treatment regimens delivered through once daily oral gavage: vehicle, dabrafenib + trametinib (one of the current clinical therapeutic regimens for BRAF^V600E^+ lung cancer patients; D+T), and afatinib (pan-ERBB inhibition). Tumors from the CTG0167 model, D+T treatment only exhibited stable disease, whereas afatinib showed clear and stable tumor regression over 32 days of treatment (**Figure 3A**). Strikingly, mice implanted with CTG0167 with complete responses were taken off afatinib following the study, but still showed no tumor reoccurrence after an additional 30 day waiting period (data not shown). Mice implanted with NCI349418 exhibited responses to both D+T and afatinib, showing durable tumor regression with either therapeutic strategy (**Figure 3B**). We further characterized the responses in this exquisitely sensitive model through immunohistochemical analyses of FFPE tumor sections (Supplementary Figure 4). SPC and NKX2.1 are markers of alveolar fate identity, whereas total EGFR and pERK are markers for the potential for MAPK signaling in this model ^24^. Both D+T and afatinib treated tumors showed clear reductions in MAPK signaling compared to vehicle controls, but no obvious deviations in alveolar identity, histology, or differentiation status. Moreover, mice that harbored HCIBL1 tumors exhibited stable disease, followed by progression in response to D+T, but had a stronger response to afatinib: 3 weeks of stable tumor shrinkage followed by tumor progression (**Figure 3C**). Next, Lito7S showed stable disease followed by progression to both D+T and afatinib, but with a marginally longer delay until tumor volume increase with afatinib (**Figure 3D**). Finally, the Lito1S model, a notably aggressive and fast growing model, revealed no response to D+T, but a short response to treatment with afatinib (**Figure 3E**). Notably, treatment with afatinib significantly extended survival in this model compared to D+T by 3 weeks (**Figure 3F**; *p* < 0.0001 Log Rank Mantel Cox.) Overall, this data revealed a previously unknown role for ERBB signaling in regulating tumor maintenance in 5 different BRAF^V600E^+ PDX models.

**Figure 3.**
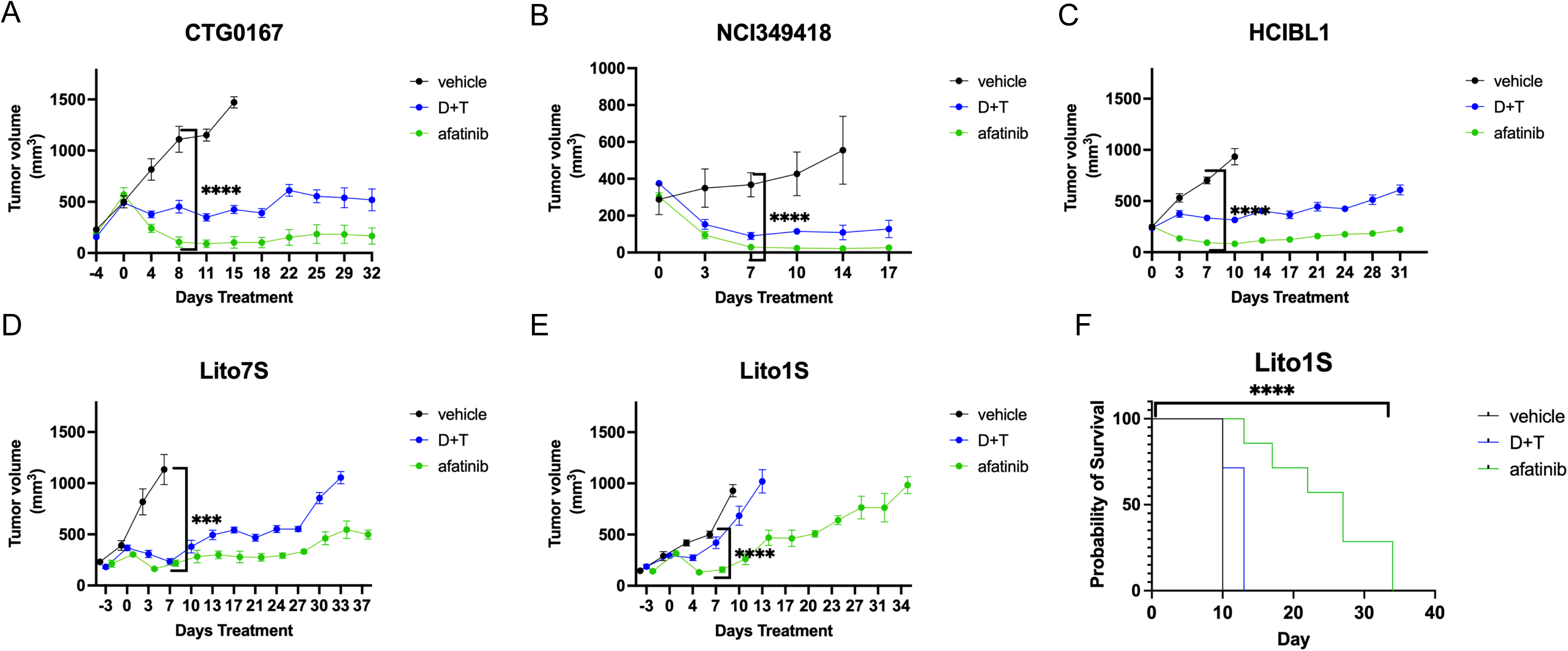
Pan-ERBB Pharmacological Inhibition Has Potent Anti-Tumor Activity Against BRAF^V600E^ PDX Models. Quantification of tumor volume in immunocompromised NOD-SCID mice implanted with A.) CTG0167 B.) NCI349418 C.) HCIBL1 D.) Lito7S E.) Lito1S BRAF^V600E^+ patient-derived xenograft PDX tumors treated once daily via oral gavage with vehicle (black), 75 mg/kg dabrafenib + 1 mg/kg trametinib (blue), 15 mg/kg afatinib (green). Error bars indicate the SEM. Statistical analysis was performed using a one-way ANOVA, where ^∗∗∗∗^p < 0.0001. n = 25-40 mice/PDX model with 5-8 per group. F.) Kaplan Meier survival curve analyses of mouse survival over 40 days of the indicated drug treatments shown in E.). Statistical analyses were performed using a log-rank Mantel-Cox where ^∗∗∗∗^p < 0.0001.

**Table 1:**
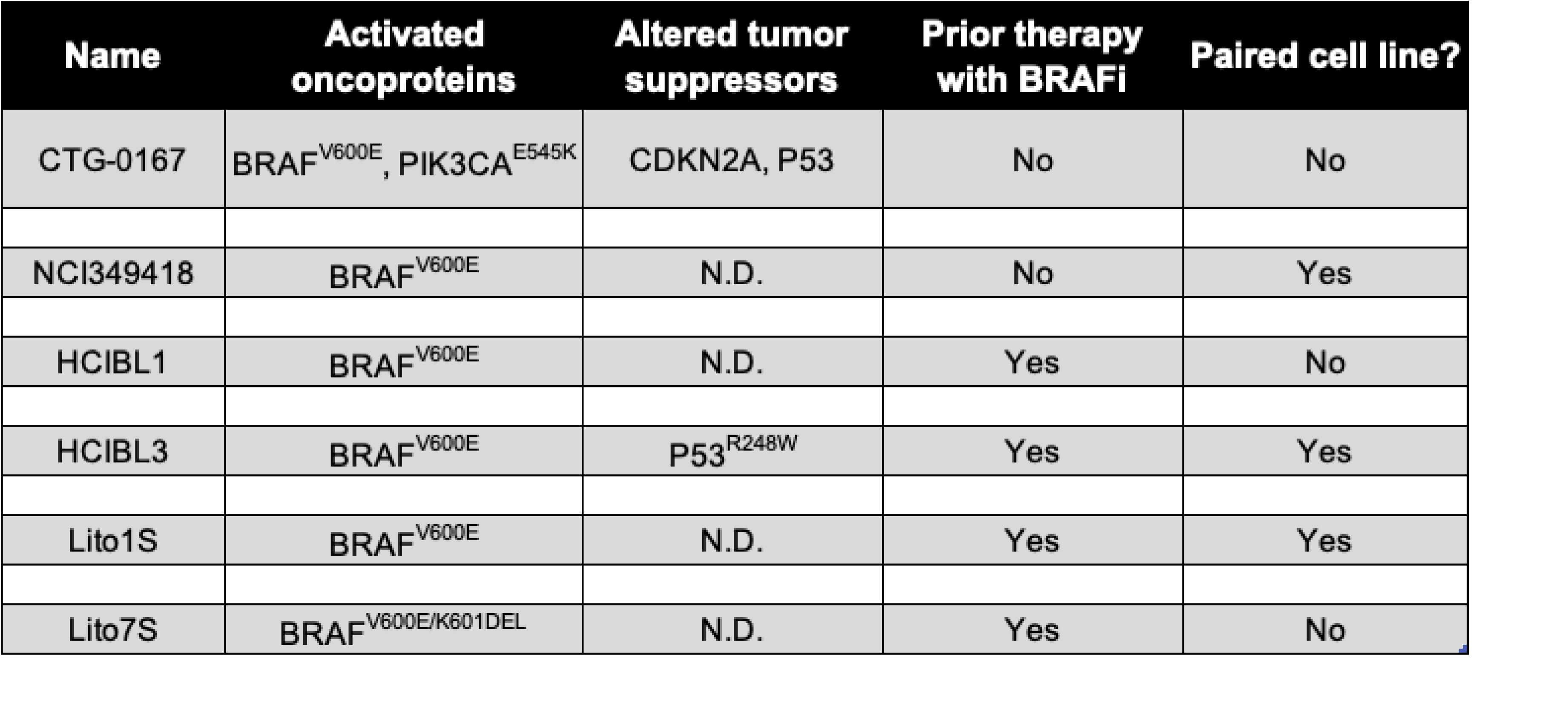
Lung Cancer BRAF^V600E^ PDX Bank.

| Name | Activated oncoproteins | Altered tumor suppressors | Prior therapy with BRAFi | Paired cell line? |
| --- | --- | --- | --- | --- |
| CTG-0167 | BRAF<sup>V600E</sup>, PIK3CA<sup>E545K</sup> | CDKN2A, P53 | No | No |
| NCI349418 | BRAF<sup>V600E</sup> | N.D. | No | Yes |
| HCIBL1 | BRAF<sup>V600E</sup> | N.D. | Yes | No |
| HCIBL3 | BRAF<sup>V600E</sup> | P53<sup>R248W</sup> | Yes | Yes |
| Lito1S | BRAF<sup>V600E</sup> | N.D. | Yes | Yes |
| Lito7S | BRAF<sup>V600E/K601DEL</sup> | N.D. | Yes | No |

We were surprised by the ability of single-agent ERBB inhibition to exhibit control over tumor volume in BRAF^V600E^+ tumors, and wanted to explore a deeper understanding of how this occurs biologically. To start, we examined the mechanism of how afatinib, a type 6 irreversible small molecular TKI, binds to EGFR. Afatinib forms an irreversible, covalent bond with Cysteine797 of EGFR. This bond can be disrupted through a point mutation of Cystine797 to a Serine (EGFR-C797S), which eliminates the active site residue required for the warhead or reaction moiety of afatinib to work ^31^. Albeit, despite this, previous work has still observed residual activity of afatinib as a non-covalent binding, general TKI against EGFR mutant lung cancer cells that also harbor a C797S mutation ^32^. Recent work has shown that select EGFR inhibitors may have efficacy on some human lung cancer BRAF mutation populations as a single-agent, which could help explain the striking responses we observed in **Figure 3** ^33^. To test the role of EGFR-C797 in the efficacy of afatinib treatment in BRAF^V600E^+ cells, we expressed wild-type (WT) or C797S EGFR in HCC364 BRAF^V600E^+ human lung cancer cells. Immunoblot analysis of cell extracts from HCC364 cells expressing an empty vector (EV) control or WT EGFR mammalian expression plasmids revealed inhibition of phosphorylated EGFR following treatment with afatinib, but not D+T (**Figure 4A**). Accordingly, downstream signaling through pERK was reduced with both D+T and afatinib, whereas pro-apoptotic BCL-2 family member BIM was activated with both inhibitors. In contrast, cell extracts expressing EGFR-C797S demonstrated biochemical resistance through phosphorylated EGFR, pERK, and rescued BIM activation in this model (**Figure 4A**). Next, we wanted to build further on this result and use this system to understand if EGFR’s role in promoting BRAF^V600E^ tumorigenesis was tumor cell autonomous or if it was mediated by the effects of EGFR in the tumor microenvironment. To that end, we implanted either EV, EGFR WT or EGFR-C797S expressing HCC364 cells in a subcutaneous xenograft model and dosed mice once daily with either vehicle (black), D+T (blue), or afatinib (green). In both EV or WT EGFR expressing xenografts, mice were sensitive to both D+T (BRAF) and afatinib (pan-ERBB) treatment strategies, where both therapeutic strategies caused durable tumor regression (**Figure 4B&C**; *p* < 0.001). However, in the HCC364 xenograft model expressing EGFR-C797S, the tumors exhibited a partial, but significant rescue of tumor growth in response to afatinib treatment (**Figure 4D&E**; *p* < 0.0001). Data from this experimental model uncovered a partially cell autonomous role for EGFR in supporting BRAF^V600E^-driven tumors, but we wondered if the partial rescue could be due to contributions from other ERBB family members, as afatinib has activity against wild-type ERBB2/HER2.

**Figure 4.**
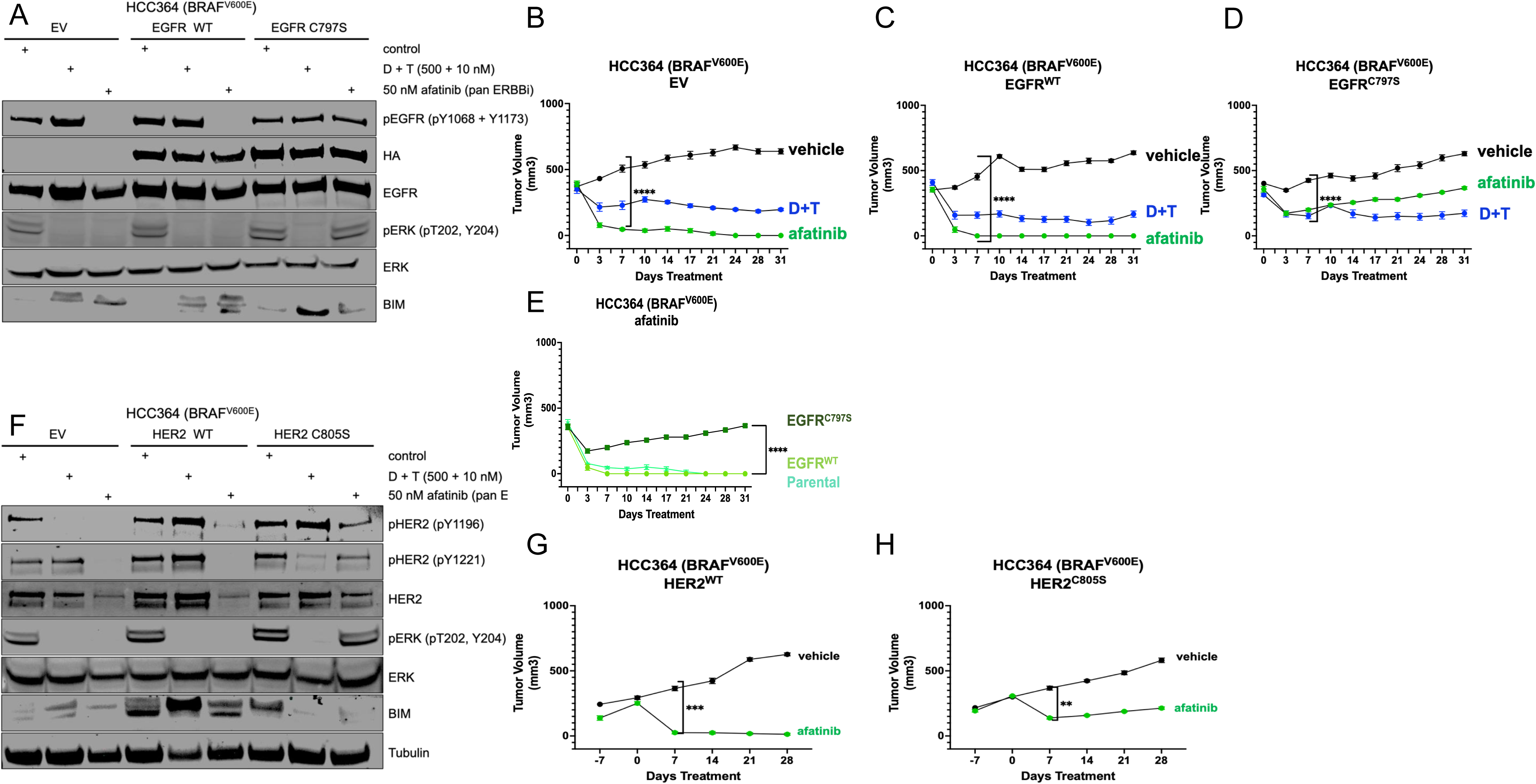
EGFR and HER2 Mediate Sensitivity to Afatinib in BRAF^V600E^ Lung Cancer Xenograft Models in a Partially Cell Autonomous Manner. A.) Immunoblot analyses of HCC364 human lung cancer cells stably expressing lentivirus encoding an empty vector control plasmid, wild-type EGFR expression plasmid, or an expression plasmid that harbors an EGFR mutation (C797S) that renders resistance to afatinib. Protein extracts were collected at the specified timepoints and probed with the indicated antibodies. B.) BRAF^V600E^+ HCC364 cell-line derived xenograft tumors stably expressing an empty vector plasmid treated once daily for 31 days via oral gavage with vehicle (black), 75 mg/kg dabrafenib + 1 mg/kg trametinib (blue), or 15 mg/kg afatinib (green). Error bars indicate the SEM. Statistical analysis was performed using a one-way ANOVA, where ^∗∗∗∗^p < 0.0001. N=5 mice per each treatment group. C.) BRAF^V600E^+ HCC364 cell-line derived xenograft tumors stably expressing a WT EGFR plasmid treated once daily for 31 days via oral gavage with vehicle (black), 75 mg/kg dabrafenib + 1 mg/kg trametinib (blue), or 15 mg/kg afatinib (green). Error bars indicate the SEM. Statistical analysis was performed using a one-way ANOVA, where ^∗∗∗∗^p < 0.0001. N=5 mice per each treatment group. D.) BRAF^V600E^+ HCC364 cell-line derived xenograft tumors stably expressing EGFR^C797S^ expression vector plasmid treated once daily for 31 days via oral gavage with vehicle (black), 75 mg/kg dabrafenib + 1 mg/kg trametinib (blue), or 15 mg/kg afatinib (green). Error bars indicate the SEM. Statistical analysis was performed using a one-way ANOVA, where ^∗∗∗∗^p < 0.0001. N=5 mice per each treatment group. E.) Direct comparison of tumor volume change in HCC364 cell line xenograft models expressing EV, WT EGFR, or EGFR^C797S^ and treated once daily for 31 days with 15 mg/kg afatinib. Error bars indicate the SEM. Statistical analysis was performed using a one-way ANOVA, where ^∗∗∗∗^p < 0.0001. N=5 mice per each treatment group. F.) Immunoblot analyses of HCC364 human lung cancer cells stably expressing lentivirus encoding an empty vector control plasmid, wild-type HER2 expression plasmid, or an expression plasmid that harbors an HER2 mutation (C805S) that renders resistance to afatinib. Protein extracts were collected at the specified timepoints and probed with the indicated antibodies. N=2. G.) HCC364 cell-line derived xenograft tumors stably expressing a WT HER2 plasmid treated once daily for 28 days via oral gavage with vehicle (black) or 15 mg/kg afatinib (green). Error bars indicate the SEM. Statistical analysis was performed using a one-way ANOVA, where ^∗∗∗^p < 0.001. N= 7 or 8 mice per group. H.) HCC364 cell-line derived xenograft tumors stably expressing a HER2^C805S^ plasmid treated once daily for 28 days via oral gavage with vehicle (black) or 15 mg/kg afatinib (green). Error bars indicate the SEM. Statistical analysis was performed using a one-way ANOVA, where ^∗∗^p < 0.01. N= 7 or 8 mice per group.

To test this new hypothesis, we engineered expression plasmids for both HER2 WT and HER2-C805S (the parallel afatinib-disabling mutation in the HER2 kinase domain) and expressed them in HCC364 cells. At the biochemical level, the EV and WT HER2 expressing cells revealed similar sensitivity to afatinib, measured by phosphorylated HER2 at two unique tyrosines (Y1196 and Y1221), with subsequent downstream effects at both pERK and BIM (**Figure 4F**). As predicted, HER2-C805S was sufficient to render resistance to afatinib, as measured by phosphoryated HER2, as well as the corresponding rescue of downstream effects through pERK and BIM (**Figure 4F**). Next, we set out to utilize these engineered cells to extend our understanding of the role(s) of HER2 in teaming up with EGFR to support BRAF^V600E^+ xenograft tumors and learn how this model responds to a pan-ERBB signaling targeted pharmacological approach. While the WT HER2 expressing HCC364 cell were still sensitive to once daily treatment with afatinib, the HER2-C805S mutant rendered a partial tumor growth rescue similar to what we observed with EGFR-C797S (**Figures 4G&H**; *p* < 0.01). We next wondered if combining the two mutants (EGFR-C797S and HER2-C805S) might be necessary to enhance the rescue effects of each single mutation. Immunoblot analyses of the two combined mutations was consistent with the individual mutation analyses, but did not reveal any clear additive or synergistic effects with inhibition of the targets (Supplementary Figure 5A). When the combined mutant cells were deployed in a xenograft model, we still saw only a partial rescue of afatinib sensitivity (Supplementary Figure 5B; *p* < 0.01), which was entirely consistent with the observations from the immunoblot analyses. Taken jointly, this data brings to light a partly tumor cell autonomous role for ERBB signaling in promoting human BRAF^V600E^+ xenograft tumor growth.

### ERBB amplifies signaling through the MAPK pathway and diversifies signaling through MTOR or c-Jun

EGFR has 12 intracellular autophosphorylation tyrosines, 10 with known signaling adaptor docking sites to multiple downstream signaling pathways ^34^. We postulated that EGFR could be contributing to signal transduction in a BRAF^V600E^+ oncogenic program by doing one of two things: further amplifying signaling through the RAS-regulated RAF>MEK>ERK pathway or diversifying signaling through the activation of additional pathways. To unbiasedly investigate these options we turned to reverse phase phosphorylation arrays or RPPA analyses. For this approach, we took human BRAF^V600E^+ Lito1S or HCC364 cells and treated them with either vehicle control (drug 1), BRAF vertical inhibition through D+T (drug 2), pan-ERBB inhibition through afatinib (drug 3), or the combination of the two pathway-blockades (drug 4) (Supplementary Figure 6A). We observed significant changes in protein expression of 221 proteins (HCC354) or 37 proteins (Lito1S), respectively (Supplementary Figure 6B&C). Indeed, these two cell lines showed a variety of unique and shared changes, but the largest effects were observed in the RAS-regulated RAF>MEK>ERK signaling cascade, including ARAF, DUSP4, and Mig6 (Supplementary Figures 6D&E). Validation of these significant protein changes through immunoblot analyses confirmed the RPPA results, including enhanced target suppression of MAPK related proteins Mig6, ARAF, phosphorylated-c-JUN at Serine 73, and phosphorylated-CDC2 at tyrosine 15, as well as other downstream pathways that regulate phosphorylated Ribosomal S6 Kinase (S6) at serines 235 and 236 (Supplementary Figure 6F). Taken as a whole, this data revealed that while EGFR may be helping diversify signaling through MTOR regulated Ribosomal S6-Kinase as well as c-JUN, it also further amplifies signal transduction through the canonical RAS>RAF>MEK>ERK MAPK cascade independent of the constitutively activated oncoprotein BRAF^V600E^.

To further scrutinize the mechanism of MAPK pathway amplification, we set out to investigate the effects of EGFR signaling on key nodes in the pathway, starting with wild-type RAS isoforms in BRAF^V600E^+ lung cancer cells. Using immunoblot analyses, we observed GTP-bound or activated RAS in BRAF^V600E+^ BP245 mouse lung cancer cells that correlated with protein expression of EGFR using a DOX-inducible shRNA model (Supplementary Figure 7A). To build on this result and broaden its importance to other driver models, we also examined whether activated RAS-GTP was affected by EGFR protein expression in EML4-ALK-driven mouse cells. Knockdown of EGFR with the same DOX-inducible shRNA system in an EML4-ALK-driven model revealed a clear reduction in RAS-GTP (Supplementary Figure 7B). This result is particularly informative because EML4-ALK can signal directly through RAS, unlike BRAF^V600E^, which is constitutively activated but a node below KRAS in the signaling cascade ^35^. Upon establishing EGFR could signal through RAS in these cells, we wanted to investigate whether a specific isoform of RAS, KRAS, was necessary for EGFR support signaling to amplify signal transduction in the canonical MAPK pathway. Here, we utilized a genetic strategy, where a dominant negative KRAS mutation, S17N, was stably introduced to the cells, and evaluated through immunoblot analyses. KRAS^S17N^ disrupts KRAS signaling by diminishing its affinity for GTP ^36^. Despite the presence of constitutively active BRAF^V600E^ in these cells, inducible activation of KRAS^S17N^ reduced pERK activity (Supplementary Figure 7C). The magnitude of this effect was further enhanced through pan-ERBB inhibition with afatinib, demonstrating ERBB signaling through KRAS was a critical contributor in this model. This outcome reveals that while BRAF^V600E^ is constitutively active, its independence from RAS is context dependent, and can receive upstream supplementation from EGFR through KRAS. Finally, we wanted to investigate the importance of MEK1/2 signaling downstream of BRAF^V600E^ in EGFR support signaling. We hypothesized that constitutive activation of MEK downstream of BRAF^V600E^, such as through MEK^S218E/S222D^ could overcome the loss of pan-ERBB blockade in BRAF-driven lung cancer cells^37^. Through immunoblot analyses using pERK as the measure for pathway activation/disruption, we noted a robust rescue of pan-ERBB blockade in BP245 cells stably expressing MEK^S218ES222D^ compared to WT MEK (Supplementary Figure 7D). The data from these experiments collectively reveals, node by node, the importance of both KRAS and MEK1/2 in mediating the support signaling by EGFR in BRAF^V600E^+ lung cancer cells.

### pEGFR is active in BRAF^V600E^+ patient samples

Next, we wanted to investigate the potential presence of activated or phosphorylated EGFR in context of BRAF^V600E^+ human lung cancer patient samples before or after treatment with different BRAF therapeutic regimens. An assorted cohort of BRAF^V600E^+ lung cancer patient samples from the MD Anderson Cancer Center were evaluated pre- or post-treatment with different clinically-relevant BRAF targeted therapy regimens. Sample 1 was pleural fluid obtained from a patient pre-treatment on a trial with vemurafenib and sorafenib for 8 months (**Figures 5A&B**). Expression of pEGFR is visible in the tumor regions through comparison with H&E staining in **Figure 5A**. Sample 2 was taken from a post treatment surgical biopsy of another patient who was treated, and responded to vemurafenib for 8 months (**Figures 5C&D**). pEGFR protein expression levels are visibly high at the time resistance was acquired, suggesting a potential role for EGFR signaling in that sample. The third and fourth samples came paired (pre-3 months treatment with D+T) from patient 3, followed by progression and the post-treatment biopsy came following a switch to treatment with chemotherapy + immunotherapy for 2 years (**Figures 5E-H**). Assessing pEGFR from the pre- to post-treatment sample shows a clear increase in pEGFR in the resistant sample (compare **Figure 5F** **to** **Figure 5H**). Samples 5 and 6 were taken pre-treatment with D+T for 10 months from patient 4, followed by 2 lines of chemotherapy and the post-biopsy (**Figure 5I-L**). While the staining for pEGFR pre-treatment appears at low levels and with a diffuse pattern, the post-treatment sample reveals a robust increase in pEGFR stain with a clear upregulation pattern that overlays with the tumor cells compared to the earlier sample (compare **Figure 5J to 5L**). Finally, for the third pair of samples we evaluated pre-and post-treatment with D+T for 5 years, followed by the identification of a KRAS^G12S^ mutation (**Figures 5M-P**) in patient 5 Again, there was a strong upregulation of pEGFR protein levels in the post-treatment resistant sample (even with the KRAS mutation detected) (compare **Figure 5N to 5P**). Interestingly, we also observed enriched, nuclear pEGFR signaling in the post-treatment sample, which have been previously linked to potentially novel or enhanced malignant functions in lung cancer models (**Figure 5P)** ^38–41^. Taken jointly, these molecular analyses showed pEGFR is present and active in a cohort of BRAF^V600E^ lung cancer patient samples, and in 3 paired samples (pre- and post-targeted therapy treatment) the post-treatment sample, higher levels of pEGFR protein correlated with the state of acquired drug resistance.

**Figure 5.**
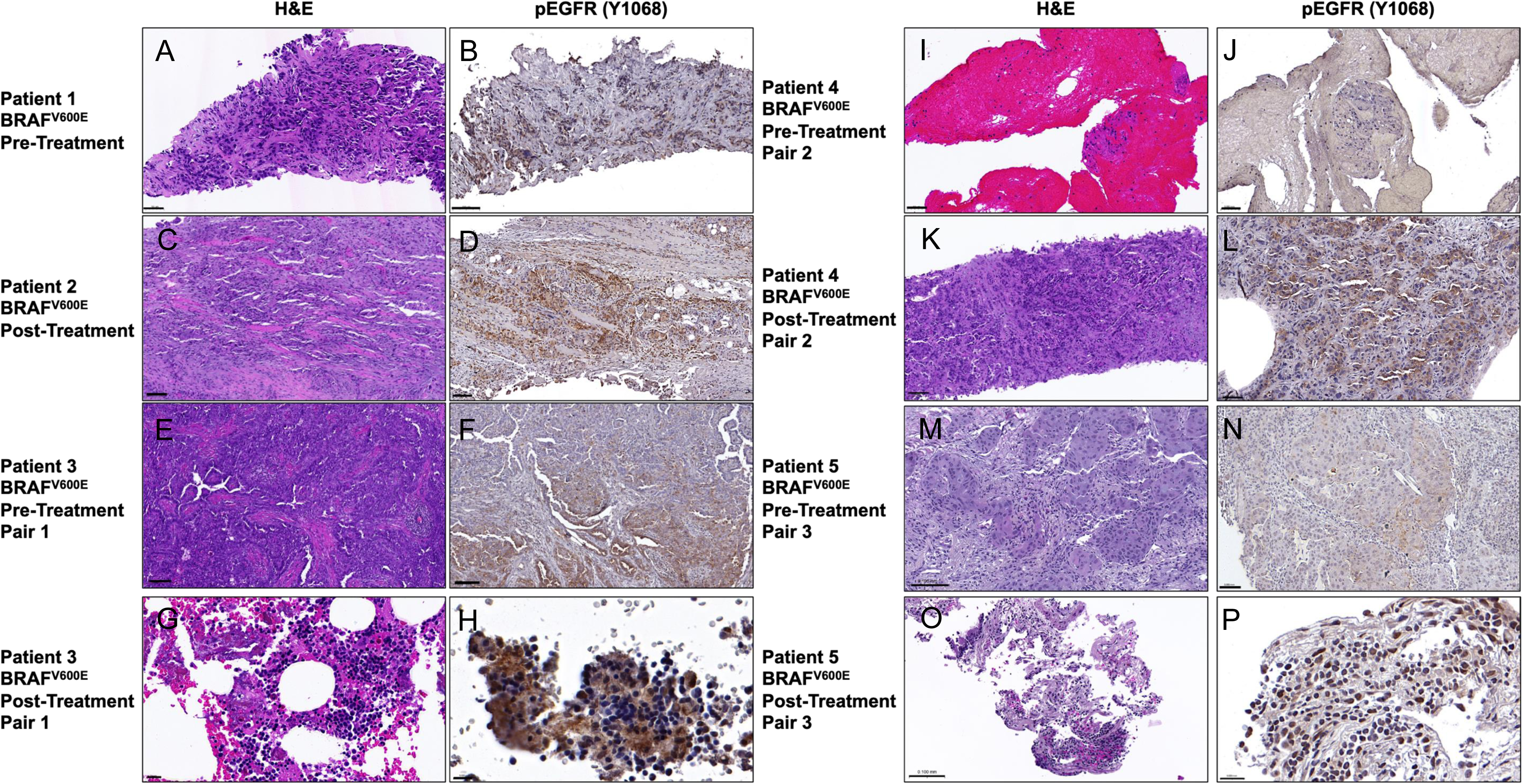
Phospho-EGFR is Activated in Human BRAF^V600E^+ Patient Samples. 8 BRAF^V600E^+ lung cancer patient tumor histological sections were stained with hemotoxilyn and eosin (H&E) or phosphorylated EGFR (Y1068), respectively. Specimens were collected either before, during, or after the patient was treated with different therapeutics. A&B.) Patient 1; FFPE from pre-treatment pleural fluid prior to treatment with vemurafenib and sorafenib on trial for approximately 8 months. 10x H&E and 20x pEGFR, respectively. C&D.) Patient 2; FFPE from post-treatment surgical biopsy (A1)—treated with and responded to Vemurafenib for 8 months prior to sample collection upon acquired resistance. 10x H&E and 10x pEGFR, respectively. E&F.) Patient 3 FFPE from pre-treatment (paired with G&H) A1 biopsy; after the patient was treated with D+T for 3 months before switching to combined one line of chemotherapy/immunotherapy. 10x H&E and 10x pEGFR, respectively. G&H.) Patient 3 FFPE from post-treatment (paired with E&F) after 2.5 years post progression on chemotherapy/immunotherapy. 40x H&E and 10x pEGFR, respectively. I&J.) Patient 4 FFPE from pre-treatment (paired with K&L) of D+T for 10 months and 2 lines of chemotherapy. 20x H&E and 10x pEGFR, respectively. K&L.) Patient 4 FFPE from post-treatment (paired with I&J) acquired resistance to treatment with D+T 10 months and 2 lines of chemotherapy. 10x H&E and 20x pEGFR, respectively. M&N.) Patient 5 FFPE from pre-treatment (paired with O&P) of D+T for 5 years. 40x H&E and 10x pEGFR, respectively. O&P.) Patient 5 FFPE during treatment (paired with M&N) with D+T for 5 years. 40x H&E and 63x pEGFR, respectively. A KRAS^G12S^ mutation was identified as a potential mediator of acquired resistance. Scale bars are shown in the bottom left corner in black.

### A Combined ERBB/BRAF therapeutic approach is superior to BRAF inhibition alone

One of the critical remaining questions was whether or not combined ERBB plus BRAF inhibition would provide more therapeutic benefit biochemically and with anti-tumor responses than targeted BRAF inhibition alone? We began by evaluating the signaling changes in HCC364 human BRAF^V600E^+ lung cancer cell extracts through immunoblot analyses following treatment with vehicle, D+T (BRAF inhibition), afatinib (pan-ERBB inhibition), or D+T+A (combined inhibition) for 2 or 24 hours, respectively. We saw clear improvement in efficacy and enhanced pathway blockade with the triple combination after 24 hours compared to 2 hours of treatment (Supplementary Figure 8A). We expanded this biochemical analyses out to a panel of new BRAF^V600E^+ primary lung cancer cells made from either PDX tumors (NCI349418 and Lito1S) or primary lung cancer patient samples (HCIBL1 and HCIBL3). Consistent with the initial observations in HCC364 cells, we saw similar superior blockade in each of the additional cell lines (Lito1S; HCIBL1, NCI349418, and HCIBL3): expanding the relevance and potency of this combinatorial approach to multiple human BRAF^V600E^ lung cancer cell line models (Supplementary Figure 8B-8E).

Next, we wanted to test the efficacy of the combination therapeutic approach *in vivo*. To that end, we utilized two BRAF^V600E^+ mouse models: the first was the autochthonous *BP245*, and the second a human-derived xenograft model, HCC364. A large cohort of *BP245* mice were again initiated with *SPC*-CRE adenovirus, and randomized onto 4 drug treatment arms: vehicle, D+T, afatinib, and the triple combination D+T+A (**Figure 6A**). Both single agent treatments resulted in a significant reduction in lung tumor burden, both quantitatively and qualitatively, but the triple combination of BRAF + pan-ERBB inhibition was clearly superior at controlling tumor burden (**Figures 6B&C**). In the HCC364 cell-line derived xenograft model, mice were initiated through subcutaneous implantation, and treated for 83 days before drug treatment was stopped (**Figures 6D&E**). Maximal tumor regression was obtained immediately (within 7 days) for single-agent afatinib or the triple combination, but took approximately 4 weeks for D+T. To resolve which agent had the deepest response on the tumors, all 3 drug treatments were removed after 83 days, and the tumor growth rebound was carefully evaluated for re-growth over the next 54 days. Consequently, the triple combination had the least tumor volumes for the reoccurring tumors, strongly reinforcing its superior treatment efficacy over both of the two single agent treatment strategies. It is important to note, however that while all tumors regrew in the D+T treated cohort, only 11/18 and 7/20 regrew in the afatinib or triple combo regimens, respectively. Thus, pan-ERBB inhibition alone and the triple combo treatment strategies had impressive “cure” rates (39% and 65%, respectively) at the time the experiment was completed. Taken collectively, this data strongly supports the potential benefit through exceptional responses when combining BRAF and pan-ERBB inhibition, up-front, in BRAF^V600E^+ lung cancers, compared to the current FDA-approved strategy of D+T alone.

**Figure 6.**
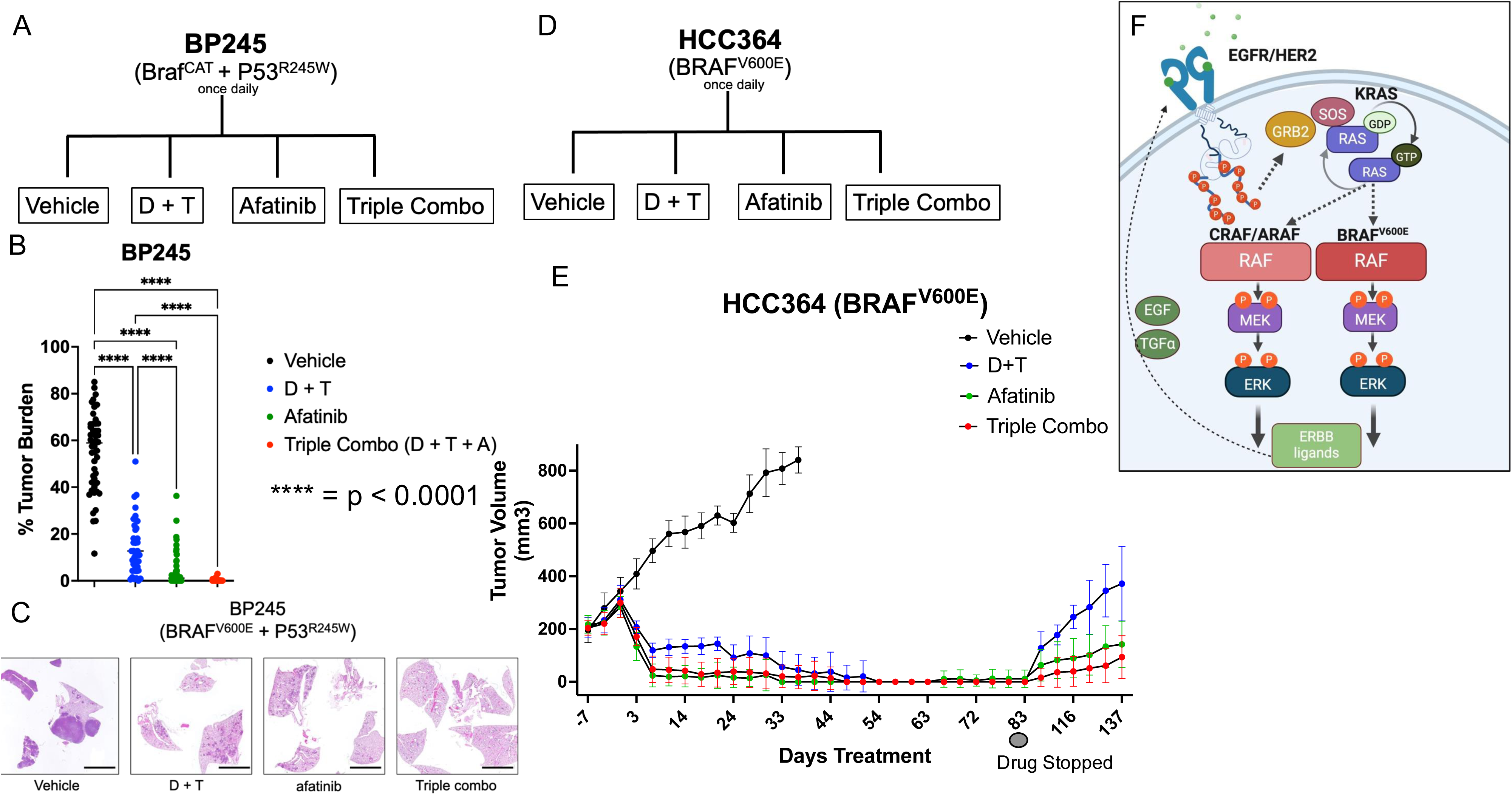
Combined EGFR and BRAF Inhibition is Superior in Durability and Depth of Response in Human and GEMM Models. A.) Schematic of the experimental design for testing EGFR and BRAF combination therapy in BP245 GEMM. B.) Lung tumor burden quantification from lungs of *BP245* mice treated for 10 weeks post initiation with vehicle, D+T, afatinib ,or the triple combination of D+T+A. Error bars indicate the SEM. Statistical analysis was performed using a one-way ANOVA, where ^∗∗**^p < 0.0001. N = 8-10 mice per group. C.) Representative images of *BP245* mouse lungs from figure 6B following treatment with the indicated therapeutic agents at euthanasia after 10 weeks of treatment post initiation. D.) Schematic of the experimental design for testing EGFR and BRAF combination therapy in HCC364 cell line derived subcutaneous xenograft model. E.) Tumor volume change over time in HCC364 cell line xenograft subcutaneous model. Flanks of immunocompromised mice were implanted with HCC364 cells and randomized to receive vehicle, D+T, afatinib, or the triple combination of D+T+A for 83 days. After 83 days drug treatment was stopped and tumors were measured out to 137 days. Error bars indicate the SEM. Statistical analysis was performed using a one-way ANOVA, where ^∗∗^**p < 0.0001. N = 10 mice per group. F.) Cartoon schematic of proposed mechanistic model summarizing results from this study: BRAF^V600E^ signaling transcriptionally upregulates ERBB family ligands (including *EGF* and *TGFalpha*.) Once activated, autocrine ERBB family signaling can activate additional signaling effectors, including RAS>CRAF and ARAF>MEK>ERK to support and promote BRAF^V600E^-driven lung cancers.

One of the most significant clinical problems with lung cancer therapies, like with all therapeutic approaches, is the inevitable path toward acquired drug resistance. As such, we wanted to explore the role of ERBB signaling in a model of acquired resistance to BRAF^V600E^-targeted therapy using the Vemurafenib Resistant, or VR1 HCC364 model ^42^. The VR1 model exhibits a phenomenon coined *oncogene overdose*, where the signaling from BRAF^V600E^ was so high, they become addicted to the reduced signaling state obtained with a BRAF inhibitor, like vemurafenib ^43^. In this model, removal of vemurafenib is cytotoxic to the cells, and as a result of that feature, they must be cultured continuously in the drug to obtain the “just-right” level of signaling in these cells (denoted by many as the “goldilocks zone”. In this set of drug-treated extracts, we evaluated the signaling changes of vehicle, D+T, afatinib, or D+T+A in VR1 resistant cells at 2 or 24 hours by immunoblot analyses. We saw enhanced pathway blockade, measured by pEGFR, pHER2, and pERK at 24 hours compared to 2 hours (Supplementary Figure 9A). To test the effects of the combination drug strategy in a resistance setting *in vivo*, we deployed a xenograft model where VR1 cells were implanted subcutaneously into immunocompromised mice. Due to the impacts of oncogene overdose in these cells, the cells had to implanted with the mouse continuously dosed on a BRAF inhibitor like vemurafenib or in this case, dabrafenib. Despite the atypical requirements of this model, we observed a significant reduction of tumor volume in the mice treated with dabrafenib + afatinib compared to afatinib alone (Supplementary Figure 9B; p = 0.0046). This data confirms the importance of EGFR support at the stage of BRAF^V600E^ acquired resistance, and demonstrates the efficacy of the combination therapeutic strategy after the evolution of resistance occurs.

The RPPA analyses in Supplementary Figure 6 revealed significant changes in other RAF family members including ARAF and CRAF, but it was not clear if those proteins were required for EGFR support signaling in BRAF^V600E^ lung cancer cell models. ARAF and CRAF can compensate for loss of BRAF through targeted inhibition and serve as by-pass signaling mechanisms in this pathway, and furthermore, have previously been shown to dimerize to promote KRAS-mutant lung cancer ^44, 45^. Finally, we wanted to explore the signaling consequences of ARAF and CRAF RNAi knockdown on BRAF^V600E^ signaling and if it mimicked the effects of pan-ERBB inhibition in our sensitive or resistant cell line models. SiRNA-mediated knockdown of either ARAF and CRAF in the HCC364 human BRAF^V600E^+ lung cancer cells led to a decrease of pERK signaling and activation of BIM (Supplementary Figure 10A). Consistent with this result, siRNA-mediated knockdown of ARAF and CRAF in VR1 resistant cells also revealed a significant reduction of pERK signaling through immunoblot analyses (Supplementary Figure 10B). These data demonstrate substantial signaling consequences of knockdown of both ARAF and CRAF in BRAF^V600E-^driven lung cancer cells pre- and post-acquired drug resistance.

## DISCUSSION

Over the last several decades our understanding of molecular subsets lung cancer defined by predictive genetic biomarkers has had a dramatic impact on the treatment of patients with biomarker-driven targeted therapies. Importantly, while BRAF^V600E^+ patients demonstrate superior responses to BRAF targeted vertical inhibition in melanoma, these responses have not been recapitulated in colorectal cancer and to a lesser degree lung cancer ^5, 46^. Indeed, in colorectal cancer, the FDA has now approved EGFR and BRAF combination therapeutic approaches, reinforcing the importance and potential benefit of EGFR inhibition in specific BRAF^V600E^+ contexts ^6, 7^.

Previous work has revealed ERBB signaling is important for AT2 cell proliferation in the normal lung, the cell of origin for lung adenocarcinoma ^47, 48^. The important role for EGFR signaling prior to lung cancer initiation may help prime its signaling to continue to be important for AT2 cells as lung adenocarcinoma initiates. Numerous previous studies have also revealed ERBB-family mediated acquired drug resistance to a variety of different oncogenic drivers, particularly in lung cancer ^49, 50^. The work we present in this study is unique from many of these past studies for evaluating the roles of ERBB signaling earlier in lung carcinogenesis prior to any therapeutic intervention. Our work is entirely consistent with previous studies in KRAS mutated lung cancer where support signaling from the ERBB pathway was identified to play a critical, supporting role ^22, 23^. This data also reinforces the importance of having tissue-specific models when evaluating mechanisms of signaling cross-talk.

Our model reveals that the initiation of constitutive BRAF signaling through the V600E mutation in the lung activates transcriptional upregulation of ERBB family ligands, including *Amphiregulin*, *HB-EGF*, and *Epiregulin* (**Figure 6F)**. Treatment with D+T reduces expression of these ligands, but their levels are not completely abrogated. Once ERBB family signaling is initiated, it can further amplify signaling through the MAPK pathway, through KRAS, CRAF/ARAF, or diversify it through activation of alternate pathways like MTOR. Pan-ERBB inhibition with afatinib can inhibit tumor maintenance as a single agent, or when combined with BRAF vertical inhibition strategies like D+T (**Figure 6F**). We were surprised by the single agent efficacy of afatinib in our models, but the pharmacological and genetic data shown here rigorously supports the notion pan-ERBB blockade significantly regulates tumor maintenance in BRAF^V600E^+ lung models. Our findings here have strong implications on our understanding BRAF^V600E^ oncoprotein signaling in different tissue specific contexts. While BRAF^V600E^ can constitutively activate signaling at this node in the pathway, our work reaffirms the notion that the RAS-independence of mutated BRAF is context-dependent in specialized tissues or microenvironments. Indeed, while BRAF^V600E^ is RAS-independent in melanoma, as expected, this is not always the case, based on our observations in the lung microenvironment ^51^.

Future studies will help identify the proper therapeutic strategy for clinical translation of BRAF^V600E^ + pan-ERBB dual blockade in lung cancer. Here, we utilized afatinib for its powerful inhibition of wild-type EGFR and pan-ERBB family blockade. Afatinib served this purpose well for our studies that were focused on mouse and human cell work, but toxicity in humans is a concern for future translational efforts. Based on the work presented here, a number of potential clinical pathway targeted strategies may prove fruitful in future efforts to find the best combination strategies to block ERBB signaling and mutated BRAF/MAPK signaling in lung cancer.

In summary, research presented here clearly indicates a clear role for ERBB signaling in supporting BRAF^V600E^-driven tumorigenesis at initiation, maintenance, and in response to therapy. Collectively, our study provide a powerful mechanistic rationale and proof of principle for the combined testing of the combination of both BRAF plus pan-ERBB inhibitors to improve the durability of patient response in cancers driven by BRAF^V600E^ mutations in lung cancer.

## AUTHOR CONTRIBUTIONS

Conceptualization: AV

Methodology/Design: AV, MM

Tools or Data: AV, MAD, MJW, MTS, BM, MN, BB, SS, CGK, WA, SP, MG, JVH, KW, and MVN

Analyses: AV, MTS, ZJ, JW

Writing: AV, MM, CGK

Funding Acquisition: AV, TB, CB, PL, MM

Supervision: AV, MM

## ACKNOWLEDGEMENTS

The authors also thank the Huntsman Cancer Institute in Salt Lake City, UT for the use of the following shared resources: (1) The Preclinical Research Resource (PRR), which provided assistance with obtaining immunocompromised mice in Utah, as well as orthoptic implantation for the experiments performed in this manuscript. (2) The University of Utah Health Science Center DNA Sequencing Core Facility. (3) The University of Utah Health Science Center Flow Cytometry Core Facility (and the National Cancer Institute through Award Number 5P30CA042014-24). We also acknowledge Novartis for providing access to dabrafenib and trametinib. We thank past members of the Vaishnavi Lab for general support of these studies including Dr. Tihui Fu, and Kyle Butler, as well as the McMahon Lab including Kayla O’Toole, Dr. Phaedra Ghazi, and Maebh Jacob, and finally the Kinsey lab including Riley Rohweder. We wish to thank the following MD Anderson Cancer Center Shared Resources: (1) The RPPA Functional Genomics Core (2) Research Histology Core Laboratory (3) DVMS Veterinary Pathology Services (4) Sanger Sequencing. The Functional Proteomics Reverse Phase Protein Array Core was supported in part by The University of Texas MD Anderson Cancer Center, P30CA016672 and R50CA221675. We wish to thank the MD Anderson Thoracic Oncology GEMINI team, including, Jeff Lewis, Waree Rinsurongkawong, Vadeerat Rinsurongkawong, Jack Lee, Jianjun Zhang, Don Gibbons, Ara Vaporciyan and John Heymach. The authors sincerely thank all of these resources for their assistance with these studies. The authors also thank BioRender for their support in creating select figures. Dr. Aria Vaishnavi is a CPRIT scholar in Cancer Research and is funded by CPRIT RR220062.

## MATERIALS AND METHODS

### Patient consent and Patient Tumor Samples

Written informed consent was obtained from the patient prior to use of the patient’s tumor samples from both the University of Utah’s Huntsman Cancer Institute and the MD Anderson Thoracic Oncology team. The consent form and protocol was reviewed and approved by the Utah Institutional Review Board.

Utah Institutional Review Board (IRB) or University of Texas MD Anderson Cancer Center IRB approval was obtained for all patients in this study. All patients provided written informed consent. Patient-Derived Xenograft Models and Human patient tissue samples were obtained from consented patient samples. The CTG0167 xenograft model was provided by Champion’s Oncology. NCI349418 was provided the National Cancer Institute Patient Derived Models Repository (PDMR). HCIBL1 and HCIBL3 (Huntsman Cancer Institute BRAF^V600E^ Lung #1 and #3, respectively) were derived at the University of Utah’s Huntsman Cancer Center under the Total Cancer Care protocol. Lito1S and Lito7S were graciously provided by Dr. Piro Lito (MSKCC).

### Vertebrate animals: breeding and experimental manipulation

All animal care and experimental procedures were approved by Institutional Animal Care and Use Committees (IACUC) at both HCI and the University of Texas MD Anderson Cancer Center. All mice were housed in environmentally controlled rooms. Mice were housed in groups in microisolator cages with bedding enrichment on ventilated racks in an AAALAC approved vivarium. Mice were provided with *ad libitum* standard chow and water through a lixit system installed into the housing racks. Mice received two health checks daily by the institute’s animal husbandry staff. Males and females were age matched within three weeks of each other, and randomly distributed equally for each experiment once they were at least 6 weeks of age. *Braf*^CAT,^ *H11^LSL-CAS^*^9^ (provided by Dr. Monte Winslow, Stanford University; RRID:IMSR_JAX:026816), *TRP53*^WM-R172H^ and *TRP53*^WM-R245W^ (provided by Dr. Guillermina Lozano, University of Texas MD Anderson Cancer Center), *C.Cg-Tg(SFTPC-rtTA)5Jaw/J* or *SPC*-rtTA (Strain #:006245; RRID:IMSR_JAX:006245), and B6;SJL-Tg(tetO-Egfr*)2-9Jek/J or *EGFR*_trunc_-tetO (Strain #: 010575; RRID:IMSR_JAX:010575). The initiation of EML4-ALK mice was performed as previously described using AdenoEA or Adeno Cas9 adenoviruses from Viraquest with permission from Dr. Andrea Ventura (MSKCC). Mice were bred as appropriate and genotyped as previously described ^24, 25^. Mouse health was assessed using the Ullmann-Cullere Body Conditioning Score (BCS) to determine whether euthanasia endpoints were met (24), at which point, mouse lungs were inflated using either PBS or 10% neutral buffered formalin for perfusion through the larynx, followed by an additional cardiac perfusion of the lung though the right ventricle of the heart until the lungs turned white. Lungs were fixed for 24 hours in formalin before transfer to ethanol for paraffin-embedding and the generation of 4 μm sections.

### Mouse experiments

Animal care and procedures were approved by the Institutional Animal Care and Use Committee Office (IACUC) of the University of Utah under protocol #18-10003 or the University of Texas MD Anderson Cancer center under protocol #00002314. For subcutaneous implantation, 10^6^ cells were resuspended into Matrigel and media and injected into each flank. For cell-line derived xenograft (CDX) subcutaneous tumorigenesis experiments, once tumors reached an average size of ≥ 250 mm^3^, mice were randomized into treatment groups based on establishing equal tumor size per group. For patient-derived xenograft (PDX) experiments, small solid tumor fragments were implanted subcutaneously into the flanks of immune-compromised mice: either of *NOD.Cg-Prkdcscid/J* or (NOD-SCID) or *NOD.Cg-Rag1tm1Mom Il2rgtm1Wjl/SzJ* or (NRG). Tumor size was measured twice weekly using digital calipers and tumor volume was calculated with the ellipsoid formula: (length x width2/2) (Tomayko and Reynolds, 1989). All dosing regimens (daily and BID) were performed 7 days a week for the period of time indicated. Dabrafenib and Trametinib were initially generously provided by Novartis. Cobimetinib was generously provided by Genentech Inc. Afatinib Dimaleate was purchased from MedChemExpress. Drugs were dissolved in corn oil or water as the vehicle and delivered by oral gavage once daily. Doxycycline or DOX chow was obtained from Inotiv (Teklad; 625 mg/kg). In conjunction with IACUC policy, mice were euthanized when tumor volume exceeded 2 cm^3^ (at the latest) or if the animal demonstrated any serious health concerns or signs of suffering. The Ullmann-Cullere Body Conditioning Score (BCS) was used to determine animal health status and to determine when and if appropriate euthanasia endpoints were met ^52^.

### Quantification and Statistical Analysis

Statistical parameters are reported in the figures and figure legends. Data are considered significant if p < 0.05. Data are presented as mean ± SEM for biological replicates. Data was analyzed using a Student’s t-test when comparing two conditions. One-way ANOVA test was performed on comparisons of more than two conditions. Survival analysis was performed using Log-rank (Mantel-Cox) test. Statistical analyses were carried out in GraphPad Prism.

### Plasmid cloning, lentivirus production, cell transduction

Recombinant adeno- or lentiviruses were administered to mice in a BSL2+ room per IACUC protocol and Institutional Biosafety Committee guidelines. Adenoviruses encoding CRE recombinase (Viraquest or the University of Iowa Viral Vector Core) were delivered through intranasal instillation using 2.5 x 10^7^ pfu, whereas the lentivirus encoding CRE (described below) was delivered through intratracheal instillation under isoflurane anesthesia with a dose 1 x 10^5^ pfu (25). Tumor initiation was performed blinded to genotype. All mice were randomized equally among experimental groups based on gender, age, and the correct genotype. All mice used in these experiments had never undergone other experimental procedures. Mice were on a mixed background of C57BL/6, 129, and FVB.

The following plasmids with either obtained or cloned/modified in the lab as described: pBabe-Puro-EGFR WT was a gift from Matthew Meyerson (Addgene plasmid # 11011 ; http://n2t.net/addgene:11011 ; RRID:Addgene_11011); pBABE-puro-ERBB2 was a gift from Matthew Meyerson (Addgene plasmid # 40978 ; http://n2t.net/addgene:40978 ; RRID:Addgene_40978); pMCL-HA-MAPKK1-WT was a gift from Natalie Ahn (Addgene plasmid # 40808 ; http://n2t.net/addgene:40808 ; RRID:Addgene_40808); pMCL-HA-MAPKK1-11A55B [S218E/S222D] was a gift from Natalie Ahn (Addgene plasmid # 40809; http://n2t.net/addgene:40809 ; RRID:Addgene_40809; pCMV-VSV-G was a gift from Bob Weinberg (Addgene plasmid # 8454 ; http://n2t.net/addgene:8454 ; RRID:Addgene_8454); psPAX2 was a gift from Didier Trono (Addgene plasmid # 12260 ; http://n2t.net/addgene:12260; RRID:Addgene_12260); lentiCRISPR v2 was a gift from Feng Zhang (Addgene plasmid # 52961 ; http://n2t.net/addgene:52961 ; RRID:Addgene_52961).

The following plasmids were designed and purchased from VectorBuilder: pLV[Exp]BsdTRE>HA/HA/HA/hKRAS[NM_033360.4]; pLV[Exp]BsdTRE>HA/HA/HA/hKRAS[NM_033360.4]*(S17N); pLV[Exp]-CMV>tTS/rtTA/Hygro; pMMLV[Exp]-hHER2_3768bp/3xFLAG-hPGK>Bsd

All site directed mutagenesis was performed using the Agilent QuikChange II XL Site-Directed Mutagenesis Kit (catalog #200522).

All miR-E doxycycline inducible lentiviral shRNA plasmids were modified/cloned from Fellman et al 2013 ^30^. shREN (control; Renilla;) shEGFR 1), 2), and 3) the following 97-mer oligos were used: 1)TGCTGTTGACAGTGAGCGACACGAGAACTAGAAATTCTAATAGTGAAGCCACAGATGTATTAGAATTTCTAGTTCTCGTGGTGCCTACTGCCTCGGA;

2)TGCTGTTGACAGTGAGCGCACCGAAATTTGTGCTACGCAATAGTGAAGCCACAGATGTATTGCGTAGCACAAATTTCGGTTTGCCTACTGCCTCGGA; 3)TGCTGTTGACAGTGAGCGACCACGAGAACTAGAAATTCTATAGTGAAGCCACAGATGTATAGAATTTCTAGTTCTCGTGGGTGCCTACTGCCTCGGA;)

Loss-of-Function CRISPR against EGFR tested 6 sgRNA that target mouse EGFR and was performed as previously described both *in vitro* for sgRNA validation and *in vivo* experiments^24^. sgRNA#2 was most effective *in vitro* and used for *in vivo* experiments. sgEGFR

1. CGGTCAGAGATGCGACCCTCGTTTTAGAGCTAGAAATAGCAAG
2. *ACTGCCCATGCGGAACTTACGTTTTAGAGCTAGAAATAGCAAG*
3. CGCGCTTACAACTGCTCGGAGTTTTAGAGCTAGAAATAGCAAG
4. TGAATCGCACAGCACCAATCGTTTTAGAGCTAGAAATAGCAAG
5. CCTCATTGCCCTCAACACCGGTTTTAGAGCTAGAAATAGCAAG
6. CTGACCGCGCTCTGCGCCGCGTTTTAGAGCTAGAAATAGCAAG

Loss-of-Function RNAi with siRNA was performed with the following reagents obtained from Horizon Discovery: ON-TARGETplus Human ARAF (369) siRNA - Individual, 5 nmol (catalog# J-003563-09-0005), ON-TARGETplus Human RAF1 (5894) siRNA - Individual, 5 nmol (catalog # J-003601-13-0005), or ON-TARGETplus Non-targeting Control siRNAs (catalog #D-001810-01-05). Each siRNA was resuspended and stored in a 10 μM stock and used at a final concentration of 25 nM.

### Cell lines, 2D and 3D culture conditions, and imaging

#### Cultured cell lines

The following cell lines were used from the following sources: HCC364 and HCC364 Vemurafenib Resistant 1 or VR1 (a gift from Dr. Trevor Bivona; UCSF), EAV (a gift from Dr. Andrea Ventura; MSKCC), EA1M (a gift from Dr. Raphael Nemenoff; University of Colorado Anschutz Medical Center) Phoenix AMPHO (ATCC; CRL 3213) and HEK-293T (ATCC; CRL 3216). All cell lines were cultured in RPMI (Roswell Park Memorial Institute) 1640 media supplemented with 10% fetal bovine serum (FBS), 1% penicillin plus streptomycin and cultured at 37°C in an atmosphere of 5% CO_2_. This will herein be referred to as 10% RPMI 1640. VR1 HCC364 cells were continuously cultured in 10 μM vemurafenib. HEK-293T and Phoenix AMPHO cells were maintained in DMEM media supplemented with 10%(v/v) FBS and 1% penicillin plus streptomycin. All established human cell lines used for these studies have been authenticated by STR profiling and Mycoplasma testing is done quarterly using PlasmoTest (InvivoGen; rep-pt1). The following cell lines were derived from the corresponding PDX tumor models: NCI349418, HCIBL1 and HCIBL3.

PDX-or human lung tissue sample derived lung cell lines were established by dissociating lung tumor tissues minced with a razor and scissors in digestive media comprised of collagenase (400 U/mL; Life Tech #17100–017), dispase (5 U/mL; Corning # 354235), elastase (4 U/mL; Worthington 2279), and DNaseI (0.25 mg/mL; Sigma DN25–100 mg) in advanced DMEM:F12 HAM media in a 37°C shaker for 30 minutes. The resulting single-cell suspension was strained using 100, 70, and 40-μm filters. Red blood cell (RBC) lysis was performed at room temperature by incubating each sample with 1x RBC Lysis Buffer (eBioscience; 00–4333–57). Cells were cultured in different ratios of Airway Epithelial Cell Basal Medium + Epithelial Cell Growth Kit (ATCC’ PCS-300-040 and PCS-300-030) and later with 5% DMEM.

### IHC and immunofluorescence of lung sections

IHC was performed as previously described (32), with the following reagents: Xylene (Thermo Fisher Scientific; UN1307), Antigen Retrieval: Citrate Buffer pH6 (Sigma-Aldrich; #C9999), Peroxide Block: BLOXALL Blocking Solution (Vector Laboratories; SP-6000–100), Protein Block: Normal Horse Serum Blocking Solution 2.5 (Vector Laboratories; S2012–50); and primary antibodies: pro-SPC (Millipore; AB3786; 1:2,000), NKX2.1/TTF-1 (Abcam; 76013-EP1584Y; 1:250; RRID:AB_1310784), EGFR-pY1068 (clone D7A5) XP (Cell Signaling Technology; 3777; 1:200), EGFR-pY1068 (Biocare Medical; API 300 AA); pERK T202/Y204 D13.14.4E XP (Cell Signaling Technology; 4370; 1:600; RRID: AB_10694057); Secondary antibody: ImmPRESS horse anti-rabbit IgG polymer kit; Peroxidase (Vector Laboratories; MP7401), DAB: ImmPACT DAB Eqv Peroxidase (horseradish peroxidase) Substrate (Vector Laboratories; SK-4103–400), Harris Hematoxylin Solution: (Sigma; HHS32), Acid Alcohol (Thermo Fisher Scientific; 6769008), Bluing solution: Scott’s Tap Water 26070–07 (VWR; 100504–452) and mounted with Permount Mounting Medium (Thermo Fisher Scientific; SP15– 500). Similarly, immunofluorescence staining of the SB SB11 transposase was performed by fixing mouse lungs in zinc-buffered formalin, processed, and embedded in paraffin, cutoff into 5-μm sections, and mounted on glass slides. Citrate-mediated antigen retrieval was performed, followed by staining with the indicated primary antibody (R&D Systems; #AF2798).

### Slide scanning, imaging, and histological analyses and quantification

Following H&E staining of sectioned lungs from remaining figures, as well as IHC, or immunofluorescence analysis, slides of sectioned mouse lungs from each indicated genotype were loaded and scanned automatically using a 3D Histech Pannoramic MIDI scanner (Thermo Fisher Scientific). Slides were imaged and analyzed using CaseViewer Software or the QuantCenter analytical center provided by 3D Histech, and experimental identifiers were blinded for all histological and immunohistochemical analyses. Tumor burden was manually calculated on each lung lobe and total tumor area was compared with total lung area. Tumor diameters were measured using QuantCenter software from 3D Histech. Cellprofiler was used to quantitate median fluorescence intensity with a previously described pipeline following immunofluorescence analysis of mouse tumor-bearing lungs (13).

### Immunoblotting/Pull-downs

Cells were treated with drugs at the indicated concentrations and for the time indicated as outlined in each figure. Protein extracts were harvested by lysing cells in RIPA lysis buffer with Halt Protease and Phosphatase Inhibitor Cocktail (Thermo Scientific) and diluted into NuPAGE LDS (lithium dodecyl sulfate, pH 8.4) sample loading buffer (Invitrogen) prior to boiling for 10 minutes at 95°C. For lysates made from tumors, mice were euthanized, tumors were excised, and snap frozen in 70% ethanol and dry ice for storage at −80°C. Later, tumors were homogenized with a glass tissue homogenizer in RIPA buffer, and a BCA assay was performed to determine protein concentration prior to boiling. Proteins were separated using standard SDS-PAGE gel electrophoresis with 4%–12% gradient Bis-Tris polyacrylamide gels, transferred to PVDF membranes for immunoblot analysis using an iBlot2 (Invitrogen), and stained with indicated primary antibodies as indicated in each figure. LI-COR brand Intercept (TBS) blocking buffer and IRDye secondary antibodies 680RD and 800CW were utilized for all immunostaining experiments. Membranes were scanned and analyzed using the Odyssey Imaging System and software (LI-COR). The primary antibodies used for immunoblot analysis are described in the key resource table and listed here:. Anti-pT202/pY204-ERK1/2 (D.13.14.4E); Cell Signaling; (Catalog# 437OS; RRID: AB_10694057). Anti-total ERK( L34F12); Cell Signaling; Catalog t# 4696S; RRID: AB_10694988). Anti-pY1068/pY1173-EGFR (1H12/2236S); Cell Signaling (Catalog# 53A5/4407S; RRID: AB_331795). Anti-total EGFR (D38B1); Cell Signaling (Catalog# 4267S; RRID: AB_2246311). Anti-pS473-AKT (D9E); Cell Signaling ((Catalog #4060L; RRID: AB_2315049). Anti-total AKT (40D4); Cell Signaling (Catalog# 2920S; RRID: AB_1147620); Anti-HB EGF (E5L5T); Cell Signaling (Catalog # 27450): Anti-EGF (F7W4J); Cell Signaling (Catalog # 82838) : Anti-BIM (C34C5); Cell Signaling (Catalog # 2933): Anti-GFP (D5.1); Cell Signaling (Catalog # 2956); Anti-β-TUBULIN (9F3): Cell Signaling (Catalog # 2128); anti-HA tag (6E2) Cell Signaling (Catalog #2367); anti-FLAG (M2); Millipore Sigma (Catalog #F1804); Anti-pHER2 Y1121/1222 (6B12) & Y1196 (D66B7); Cell Signaling (Catalog # 2243 & # 6942); Anti-HER2 (XP D8F12): Cell Signaling (Catalog # 4290); Anti-MIG6: Cell Signaling (Catalog #2440); Anti-ARAF(D2P9P); Cell Signaling (Catalog # 75804 & 4432); Anti-CRAF (D5X6R); Cell Signaling (Catalog # 12552); Anti-DUSP4 (D9A5); Cell Signaling (Catalog #5149); Anti-pS6 S235/S236; Cell Signaling (Catalog # 2211); Anti-pCDC2 Y15 (10A11); Cell Signaling (Catalog #4539); Anti-p cJUN pS73 (D47G9 XP); Cell Signaling (Catalog # 3270); Anti-RAS; Cell Signaling (Catalog #3965); Anti-pMEK (41G9); Cell Signaling (Catalog #9154). Anti-MEK (L38C12); Cell Signaling (Catalog # 4694). N = 3 replicates for each immunoblot shown.

Activated RAS pulldowns were performed with the pan-RAS activation assay from Cell Bio Labs (#STA-400) using the manufacturer’s protocol followed by immunoblot analyses. Heparin bead pull-downs were performed using Heparin Agarose Beads, followed by immunoblot analyses.

### RPPA analyses

Cellular proteins were denatured in a 1% SDS + 2-mercaptoethanol buffer solution and diluted in five 2fold serial dilutions in dilution lysis buffer. Serially diluted lysates were arrayed on nitrocellulose-coated slides (Grace Bio-Labs) by the Quanterix (Aushon) 2470 Arrayer (Quanterix Corporation). A total of 5808 spots were arrayed on each slide including spots corresponding to serially diluted (1) standard lysates, and (2) positive and negative controls prepared from mixed cell lysates or dilution buffer, respectively. Each slide was probed with a validated primary antibody plus a biotin-conjugated secondary antibody. Antibody validation for RPPA is described in the RPPA Core website: https://www.mdanderson.org/research/research-resources/core-facilities/functional-proteomics-rppacore/antibody-information-and-protocols.html. Signal detection was amplified using an

Agilent GenPoint staining platform (Agilent Technologies) and visualized by DAB colorimetric reaction. The slides were scanned (Huron TissueScope, Huron Digital Pathology) and quantified using customized software (Array-Pro Analyzer, Media Cybernetics) to generate spot intensity. Relative protein level for each sample was determined by RPPA SPACE (developed by MD Anderson Department of Bioinformatics and Computational Biology, https://bioinformatics.mdanderson.org/public-software/rppaspace/) ^53^ by which each dilution curve was fitted with a logistic model. RPPA SPACE fits a single curve using all the samples (i.e., dilution series) on a slide with the signal intensity as the response variable and the dilution steps as the independent variable. The fitted curve is plotted with the signal intensities, both observed and fitted, on the y-axis and the log2 concentration of proteins on the x-axis for diagnostic purposes. The protein concentrations of each set of slides were then normalized for protein loading. Correction factor was calculated by (1) median-centering across samples of all antibody experiments; and (2) median centering across antibodies for each sample. Results were then normalized across RPPA sets by replicates-based normalization as described ^54^. Details of the RPPA platform as performed by the RPPA Core were previously described ^55^.

## Supplementary Figure Legends

Supplementary Figure 1. Targeted BRAF^V600E^ Inhibition Diminishes Expression of HB-EGF in human lung cancer cells.

A). Immunoblot analyses with antibodies against the indicated molecules of Heparin Sepharose Immunoprecipitation (IP) in HCC364 human lung cancer cells by either collected conditioned media, standard lysate IP, or whole cell lysate (input) of cells with 500 nM dabrafenib with 10 nM trametinib (D+T) for 2 or 24 hours, respectively.

B.) Cartoon schematic of the *tetO*-*EGFR*^truncated^ ligand trap/dominant negative allele at the cellular level.

C.) Cartoon schematic of the molecular mechanism of the ES or *tetO*-*EGFR^-truncated^*; *SPC*-*rtTA* mouse. This tool is a tissue specific (lung alveolar type 2 pneumocytes), inducible, doxycycline-regulated ligand trap.

D.) Schematic of experimental design for CES mice in Figure 1B.

Supplementary Figure 2. Autocrine EGFR Signaling Supports EML4-ALK Lung Cancer Initiation.

A) Cartoon schematic of *EML4*-*ALK* CRISPR viral model and experimental plan.

B) Quantification of tumor burden in *EML4-ALK* or CAS9 alone virus in wild-type *C57Bl/6J mice or SPC*-rtTA-*EGFR-truncated (EB* or *ES* mice) at 10 weeks post initiation either + or – doxycycline chow. Error bars represent SEM. *** = p < 0.001 by an unpaired T-Test. N=8 mice per group.

C&D) H&E staining of FFPE lungs of ES mice initiated with Ad-EA virus at 10 weeks post initiation either –(C) or + (D) doxycycline chow.

Supplementary Figure 3. Lung Cancer Xenografts Reveal Sensitivity to the Anti-Tumor Activity of DOX-Inducible shRNAs Against Mouse EGFR at Different Stages of Tumorigenesis.

A. Immunoblot analyses assessing protein expression with the indicated antibodies in EA1M (EML4-ALK) cell lysates that stably express lentiviruses that encode short hairpin RNAs (shRNAs) against Renila (control) or EGFR treated with 1μg/ml doxycline for 0, 24, or 48 hours. N=5.

B. B.) Quantification of tumor volume changes in EA1M cells stably expressing control (shREN) or shEGFR expression plasmids subcutaneously implanted into NOD-SCID mice (cell line-derived xenograft or CDX). Mice were on or off doxycycline chow at the indicated timepoints and swapped at the indicated points. For shEGFR expressing xenografts red represents on DOX and black represents off DOX chow. Mice were also dosed with 15 mg/kg afatinib at the indicated times (colored in aqua). Error bars indicate the SEM. **** = p < 0.0001 by a paired T-Test. N=5-8 mice per group (5 for shREN, 8 for shEGFR – or + DOX).

C. Immunohistochemical analyses of phosphorylated ERK1/2 in FFPE from shREN (control) expressing or afatinib treated EA1M tumor CDX model.

Supplementary Figure 4. Immunohistochemical characterization of NCI349418 tumors in response to pan-ERBB or BRAF targeted pathway blockade.

Immunohistochemical analyses of FFPE sections of NCI349418 PDX model from Figure 3B post euthanasia stained with the indicated antibodies:

A. Vehicle (H&E stain)
B. afatinib (H&E stain)
C. (D+T) (H&E stain)
D. Vehicle (anti-SPC)
E. afatinib (anti-SPC)
F. D+T (anti-SPC)
G. Vehicle (anti-NKX2.1)
H. afatinib (anti-NKX2.1)
I. D+T (anti-NKX2.1)
J. Vehicle (anti-EGFR)
K. afatinib (anti-EGFR)
L. D+T (anti-EGFR)
M. Vehicle (anti-pERK pT202; Y204)
N. afatinib (anti-pERK pT202; Y204)
O. . • D+T (anti-pERK pT202; Y204)

All images were taken at 20x magnification using a 3D Histech MIDI slide scanner.

Supplementary Figure 5. EGFR^C797S^ + HER2^C805S^ Partially Rescue Sensitivity of BRAF^V600E^ Xenografts to Afatinib.

A. Immunoblot analyses of HCC364 human lung cancer cells stably expressing lentivirus encoding wild-type EGFR and wild-type (WT) HER2 expression plasmids, or expression plasmids that harbor both an EGFR mutation (C797S) and HER2 mutation (C805S). Protein extracts were collected at the specified timepoints and probed with the indicated antibodies. N=2.
B. BRAF^V600E^+ HCC364 cell-line derived xenograft tumors stably expressing either wild-type (WT) EGFR and WT HER2 or mutant EGFR^C797S^ + HER2^C805S^ plasmids treated once daily for 21 days via oral gavage with vehicle (black) or 15 mg/kg afatinib (green). Error bars indicate the SEM. Statistical analysis was performed using a one-way ANOVA, where ^∗^p < 0.05. N= 6-8 mice per group.

Supplementary Figure 6. RPPA Analyses of BRAF^V600E^ human Lung Cancer Cell Lines Demonstrates Diversification and Amplification of Downstream Signaling Support from EGFR.

A. Schematic of experimental set up for RPPA analyses following the indicated drug treatments on the HCC364 and Lito1S cell lines. Control=DMSO vehicle, Drug 1 = D+T (500 nM D+ 10 nM T) Drug 2= afatinib (50 nM) and Drug 3= D+T+A (same drug concentrations). Each individual sample was N=3.
B. Heatmap of significant protein expression in HCC364. The significant proteins were identified by ANOVA’s post-hoc Tukey’s test (p < 0.05).
C. Heatmap of significant proteins expression in Lito1S. The significant proteins were identified by ANOVA’s post-hoc Tukey’s test (p < 0.05).
D. Venn diagrams showing the overlap of significant proteins identified for HCC364 and Lito1S cell lines.
E. List of significant RPPA protein changes in HCC364 and Lito1S cell lines
F. Immunoblot analysis for validating RPPA results in HCC364 and Lito1S cell lines. Residual cell extracts from the RPPA analyses treated with the indicated inhibitors and probed with the indicated antibodies.

Supplementary Figure 7. RAS is Activated by EGFR in Lung Cancer Cell Lines and MEK Can Help Overcome the Sensitivity of BRAF^V600E^ Cells to Afatinib.

A. Immunoblot analyses of lysates from RAS-GTP pulldowns in BP245 cell line stably expressing shEGFR. Lysates were treated with 1 μg/ml DOX for 0 or 24 hours prior to pull-down followed by immunoblot analyses with the indicated antibodies. N=2
B. Immunoblot analyses of lysates from RAS-GTP pulldowns in EA1M cell line stably expressing shEGFR. Lysates were treated with 1 μg/ml DOX for 0, 24, or 48 hours prior to pull-down followed by immunoblot analyses with the indicated antibodies. N=2
C. Immunoblot analyses of HCC364 cells transiently expressing a DOX-inducible HA-KRAS^S17N^ dominant negative construct. Extracts were treated with 1 μg/ml DOX for 0 or 24 hours and treated with 50 nM afatinib prior to harvest and immunoblots were probed with the indicated antibodies. N=3
D. Immunoblot analyses of BP245 cells expressing either WT MEK or constitutively activated MEK^S218E/S222D^ with or without 50 nM treatment with afatinib. N=3

Supplementary Figure 8. Combined BRAF and pan-ERBB Pathway Targeted Therapies Demonstrate Superior Biochemical Response Compared to BRAF Inhibition Alone.

Immunoblot analyses of cell extracts from the indicated BRAF^V600E^ lung cancer cells following 2- or 24-hours of drug treatment with the indicated inhibitors (control, D+T, afatinib, or triple combination D+T+A) and probed with the antibodies listed. D+T represents 500 nM Dabrafenib + 10 nM trametinib, A represents 50 nM afatinib represents, and triple combo is D+T+A at the same treatment concentration doses. N=3

A. HCC364 (2- and 24-hour treatments).
B. Lito1S
C. HCI-BL1
D. NCI349418
E. HCI-BL3

Supplementary Figure 9. Combined BRAF and pan-ERBB Pathway Targeted Therapies Are Effective in a Model of BRAF^V600E^+ Lung Cancer Cell Acquired Resistance.

A. Immunoblot analyses of cell extracts from VR1 (Vemurafenib Resistant 1) HCC364 cells following 2- or 24-hours of drug treatment with the indicated inhibitors (control, D+T, afatinib, or triple combination D+T+A) and probed with the antibodies listed. D+T represents 500 nM Dabrafenib + 10 nM trametinib, A represents 50 nM afatinib represents, and triple combo is D+T+A at the same treatment concentration doses. N=3
B. Quantification of tumor volume changes over the course of 25 days in VR1 cells subcutaneously implanted into NOD-SCID mice. Mice were on 500 nM Dabrafenib at the time of implantation, but 20 mg/kg afatinib was also delivered once daily by oral gavage in the indicated treatment group. Error bars indicate the SEM. *** = p = 0.0046 by a paired T-Test. N=5 mice per group.

Supplementary Figure 10. RNAi Against ARAF and CRAF Phenocopy pan-ERBB Inhibition in BRAF^V600E^+ Lung Cancer Models by Immunoblot Analyses.

A. Immunoblot analyses of cell extracts from HCC364 cells 48 hours post-transient transfection with the siRNA labeled (siSCR control, siCRAF, or siARAF) and probed with the antibodies listed.

B. Immunoblot analyses of cell extracts from VR1 (Vemurafenib-Resistant) HCC364 cells 48 hours post-transient transfection with the siRNA labeled (siSCR control, siCRAF, or siARAF) and probed with the antibodies listed. N=5.

## Notes

**Financial Support:** This research was supported by F32CA228267, K99CA246084, R00CA246084, and CPRIT R220062 to AV. This research was also supported by CA131261-09S2 to MM.

**Conflict of interest:** Dr. Piro Lito has received grants to his Institution from Amgen, Mirati, Revolution Medicines, Boehringer Ingelheim and Virtec Pharmaceuticals. P.L. is an advisory board member of Frontier Medicines, Ikena, Biotheryx and PAQ-Tx (consulting fees and equity in each) and has received consulting fees or honoraria from Black Diamond Therapeutics, AmMax, OrbiMed, PAQ-Tx, Repare Therapeutics, Boehringer Ingelheim and Revolution Medicines. Dr. Kaiwen Wang reports Janssen Pharmaceuticals advisory board and Honoraria from MJH Life Sciences. Dr. John Heymach reports consulting fees from AbbVie, AnHeart Therapeutics, ArriVent Biopharma, AstraZeneca, BioCurity Pharmaceuticals, BioNTech AG, Blueprint Medicines, BI, Bristol-Myers Squibb, Chugai Pharmaceutical, Curio Science, Dava Oncology, Eli Lilly & Co, EMD Serono, Genentech, GlaxoSmithKline, IDEOlogy Health, Immunocore, InterVenn Biosciences, Janssen Biotech, Janssen Pharmaceuticals, Mirati Therapeutics, Moffit Cancer Center, ModeX, Nexus Health Systems, Novartis Pharmaceuticals, OncoCyte, RefleXion, Regeneron Pharmaceuticals, Roche, Sandoz Pharmaceutical, Sanofi, Spectrum Pharmaceuticals, Takeda, and uniQure, honoraria from MJH Events, and Dr. Meil Love, and travel support from 10.13039/100000043American Association for Cancer Research. Dr. Marcelo Negrao reports consulting fees from Mirati, Novartis, Sanofi, Pfizer, Merck/Merck Sharp & Dohme, Genentech, Eli Lilly and AstraZeneca, honoraria from IDEOlogy, BIO Brasil and OncLive and other financial or non-financial interest with IDEOlogy, Ashfiedl Healthacre, ApotheCom, and Dava Oncology. Dr. Martin McMahon has served on advisory boards to Genentech Discovery Oncology.

### Competing Interest Statement

Dr. Piro Lito has received grants to his Institution from Amgen, Mirati, Revolution Medicines, Boehringer Ingelheim and Virtec Pharmaceuticals. P.L. is an advisory board member of Frontier Medicines, Ikena, Biotheryx and PAQ-Tx (consulting fees and equity in each) and has received consulting fees or honoraria from Black Diamond Therapeutics, AmMax, OrbiMed, PAQ-Tx, Repare Therapeutics, Boehringer Ingelheim and Revolution Medicines. Dr. Kaiwen Wang reports Janssen Pharmaceuticals advisory board and Honoraria from MJH Life Sciences. Dr. John Heymach reports consulting fees from AbbVie, AnHeart Therapeutics, ArriVent Biopharma, AstraZeneca, BioCurity Pharmaceuticals, BioNTech AG, Blueprint Medicines, BI, Bristol-Myers Squibb, Chugai Pharmaceutical, Curio Science, Dava Oncology, Eli Lilly & Co, EMD Serono, Genentech, GlaxoSmithKline, IDEOlogy Health, Immunocore, InterVenn Biosciences, Janssen Biotech, Janssen Pharmaceuticals, Mirati Therapeutics, Moffit Cancer Center, ModeX, Nexus Health Systems, Novartis Pharmaceuticals, OncoCyte, RefleXion, Regeneron Pharmaceuticals, Roche, Sandoz Pharmaceutical, Sanofi, Spectrum Pharmaceuticals, Takeda, and uniQure, honoraria from MJH Events, and Dr. Meil Love, and travel support from 10.13039/100000043American Association for Cancer Research. Dr. Marcelo Negrao reports consulting fees from Mirati, Novartis, Sanofi, Pfizer, Merck/Merck Sharp & Dohme, Genentech, Eli Lilly and AstraZeneca, honoraria from IDEOlogy, BIO Brasil and OncLive and other financial or non-financial interest with IDEOlogy, Ashfiedl Healthacre, ApotheCom, and Dava Oncology. Dr. Martin McMahon has served on advisory boards to Genentech Discovery Oncology.

