## Supplemental Figures for "BRAF^V600E^-Driven Lung Tumorigenesis Requires Ligand-Mediated Activation of ERBB Receptor Signaling"

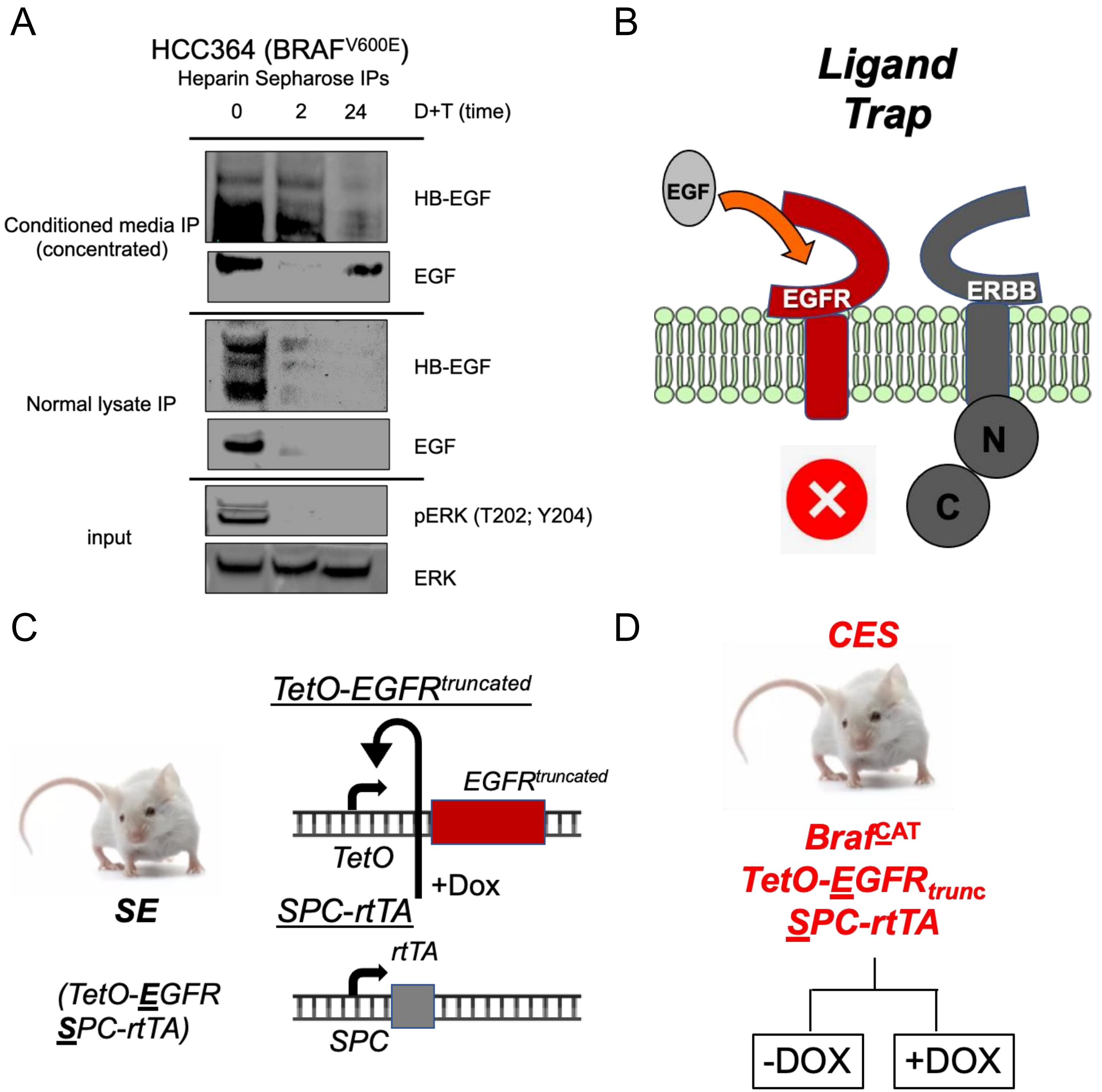


Supplementary Figure 1. ERBB Ligands are regulated by BRAF$^{V600E}$ and Contribute *In Vitro* and *In Vivo*.

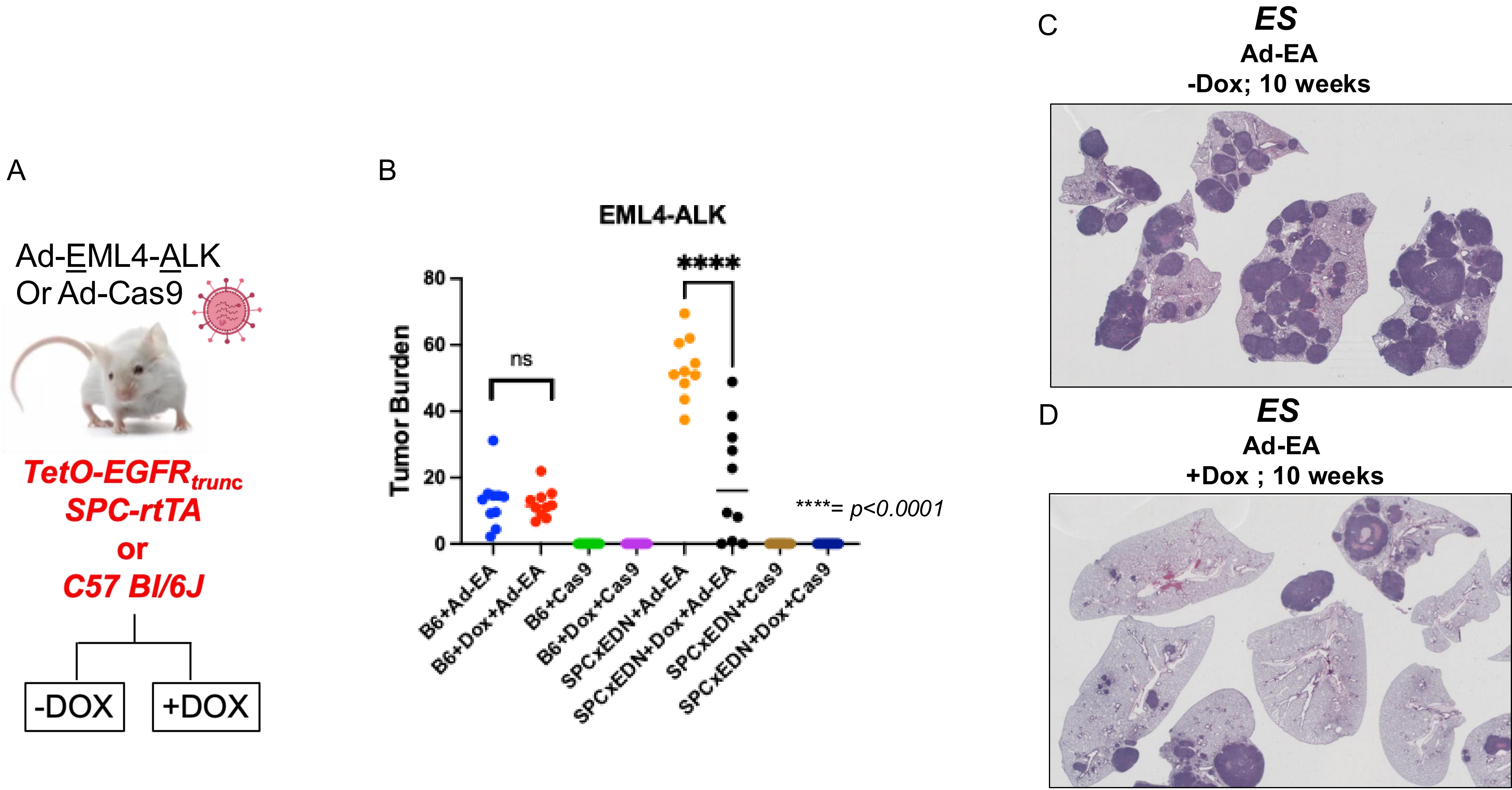


Supplementary Figure 2. Autocrine EGFR Signaling Supports EML4-ALK Lung Cancer Initiation.

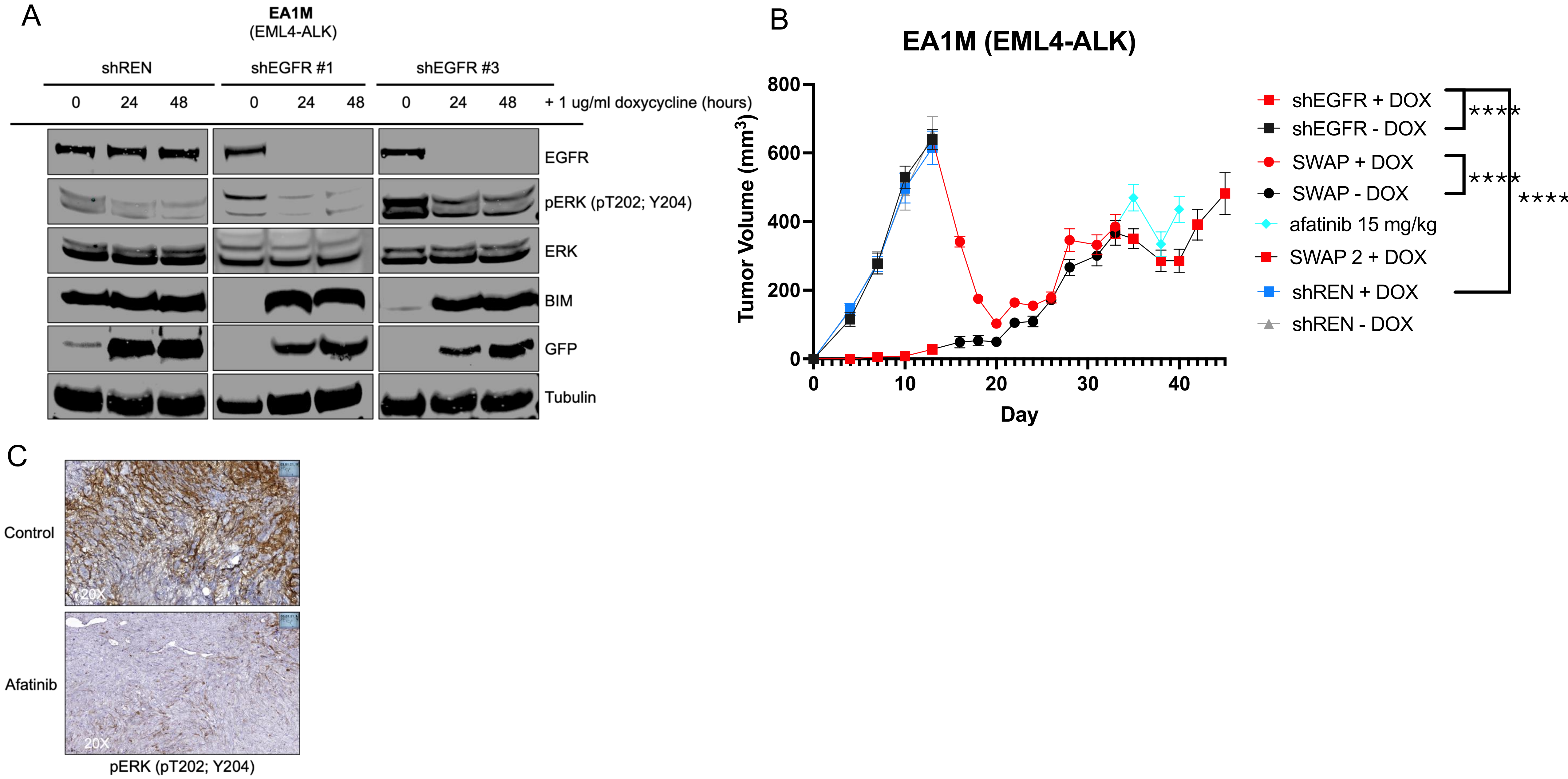


Supplementary Figure 3. Lung Cancer Xenografts Reveal Sensitivity to the Anti-Tumor Activity of DOX-Inducible shRNAs Against Mouse EGFR at Different Stages of Tumorigenesis.

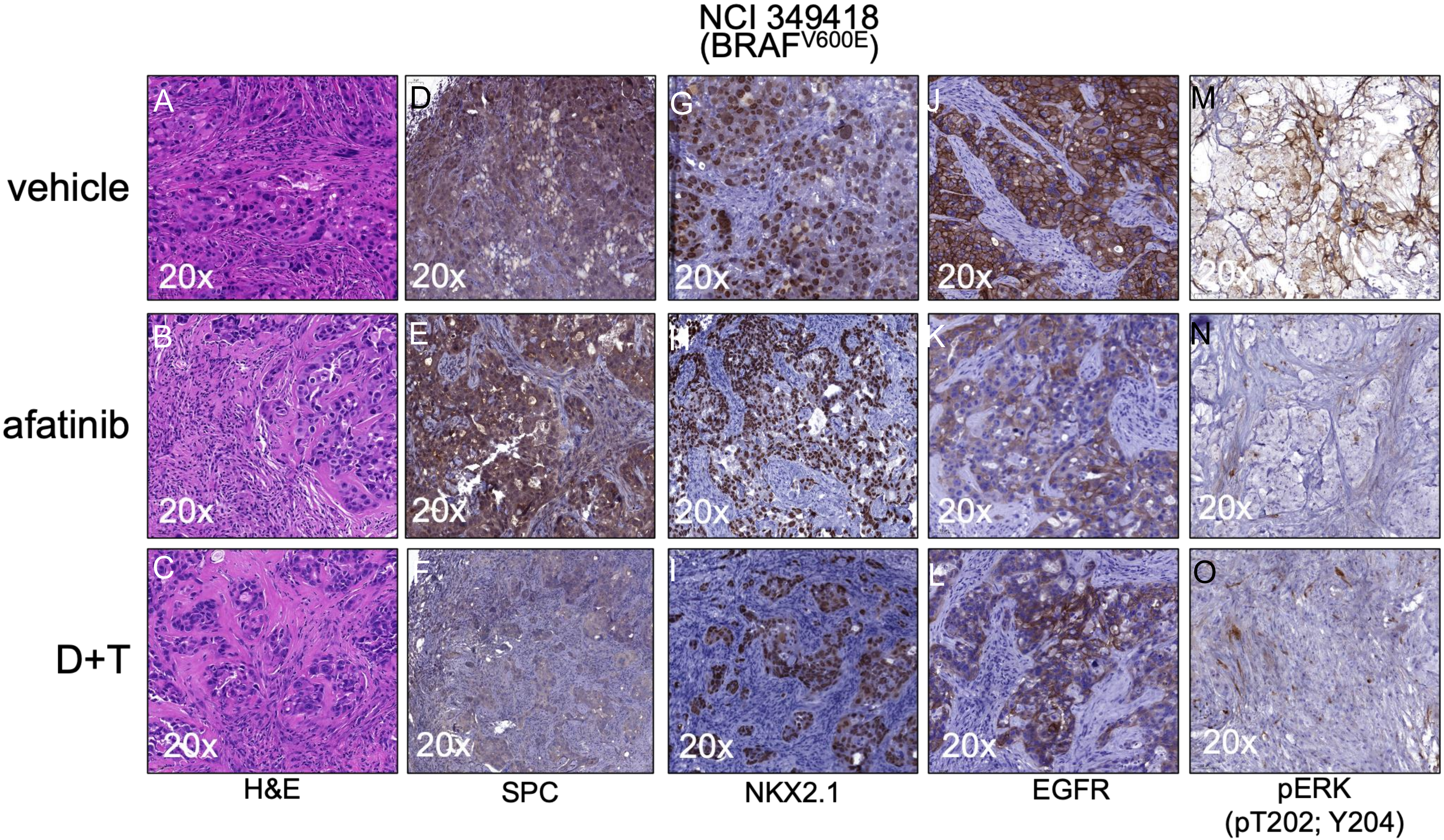


Supplementary Figure 4. Immunohistochemical characterization of NCI349418 tumors in response to pan-ERBB or BRAF targeted pathway blockade.

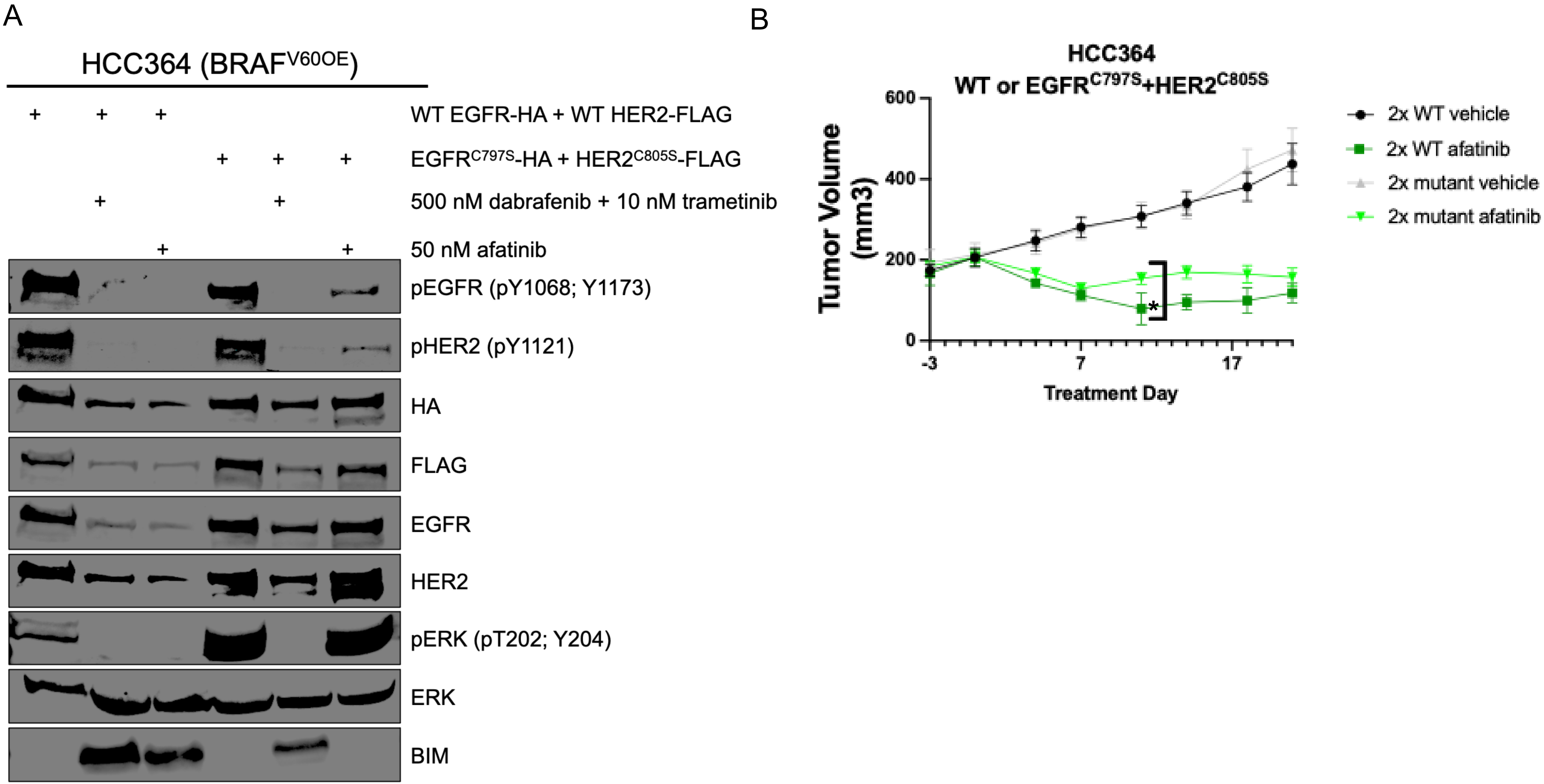


Supplementary Figure 5. $EGFR^{C797S}$ + $HER2^{C805S}$ Partially Rescue Sensitivity of $BRAF^{V600E}$ Xenografts to Afatinib.

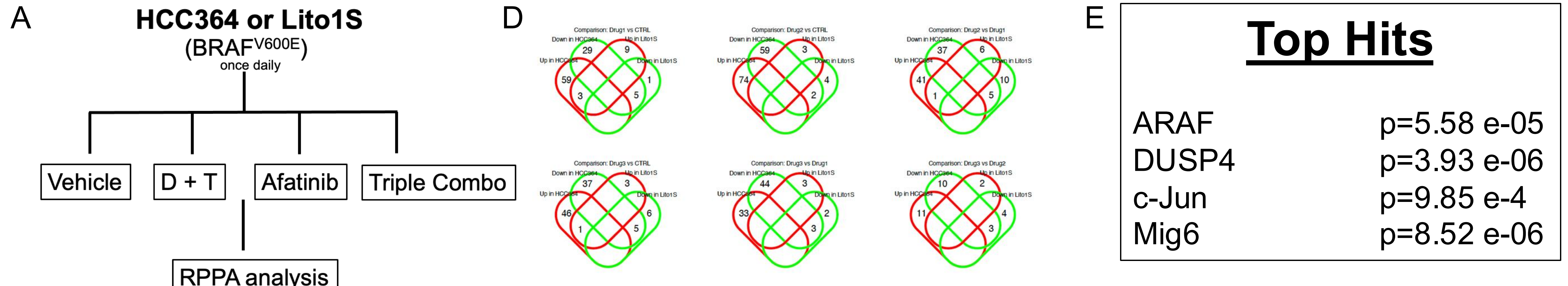

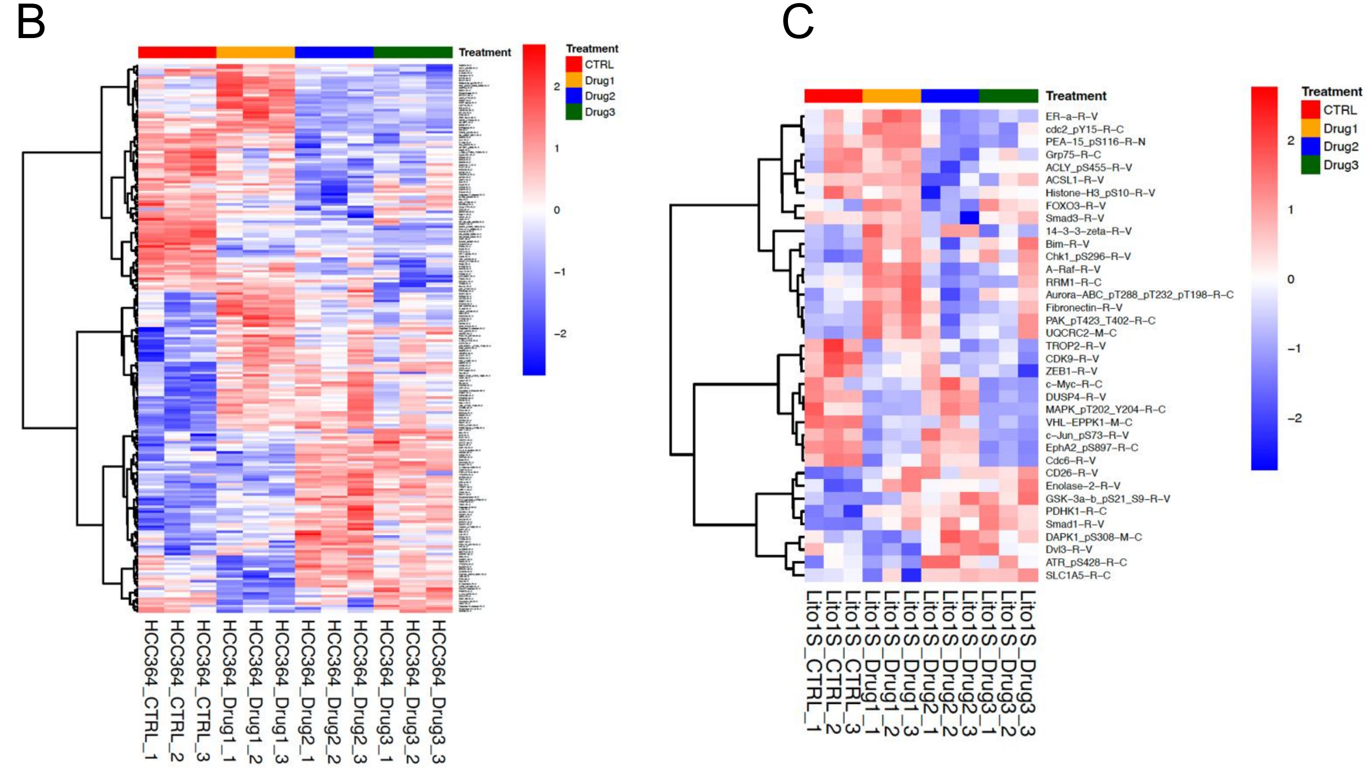

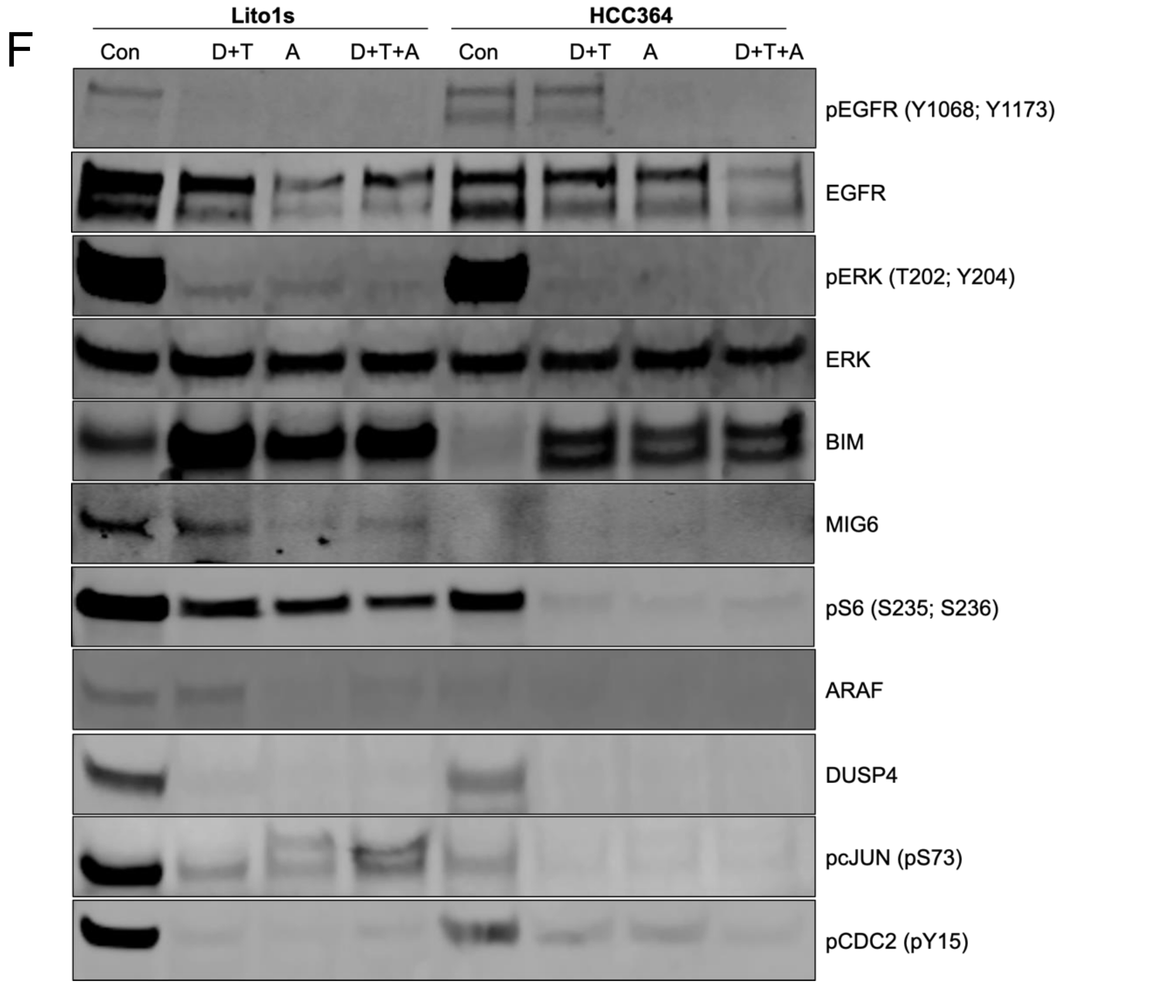


Supplementary Figure 6. RPPA Analyses of BRAF$^{V600E}$ human Lung Cancer Cell Lines Demonstrates Diversification and Amplification of Downstream Signaling Support from EGFR.

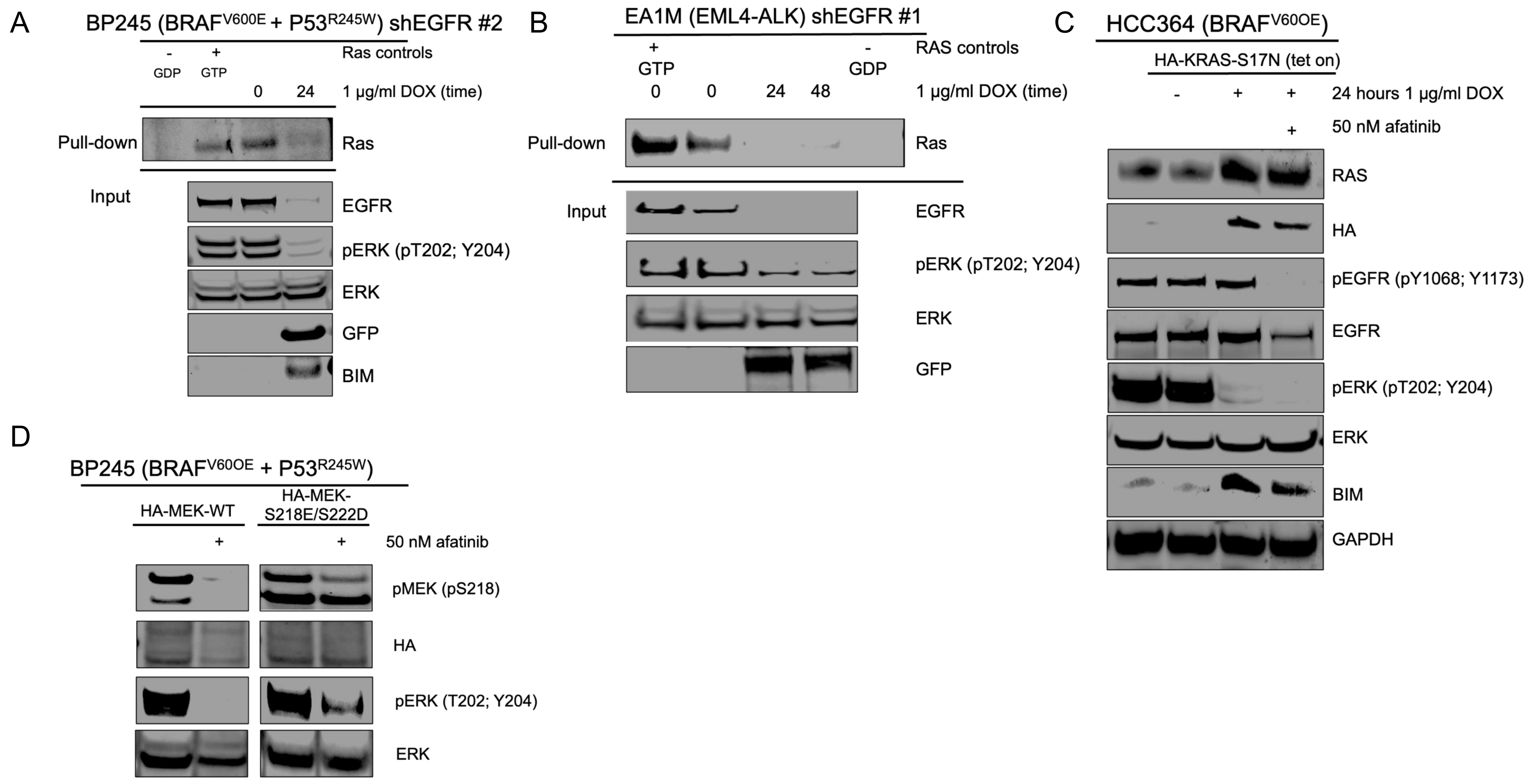


Supplementary Figure 7. RAS is Activated by EGFR in Lung Cancer Cell Lines and MEK Can Help Overcome Sensitivity of $BRAF^{V600E}$ Cells to Afatinib.

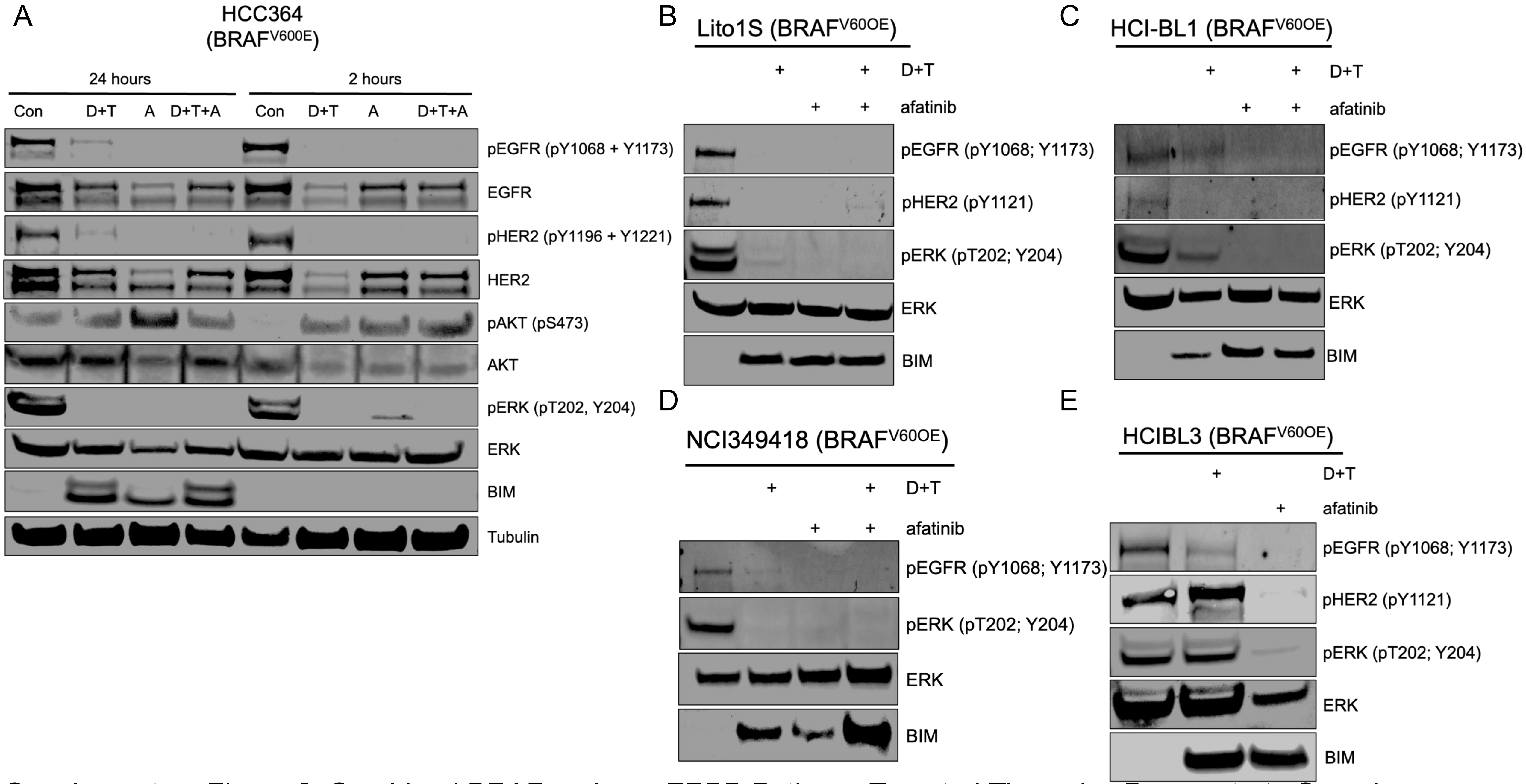


Supplementary Figure 8. Combined BRAF and pan-ERBB Pathway Targeted Therapies Demonstrate Superior Biochemical Response Compared to BRAF Inhibition Alone.

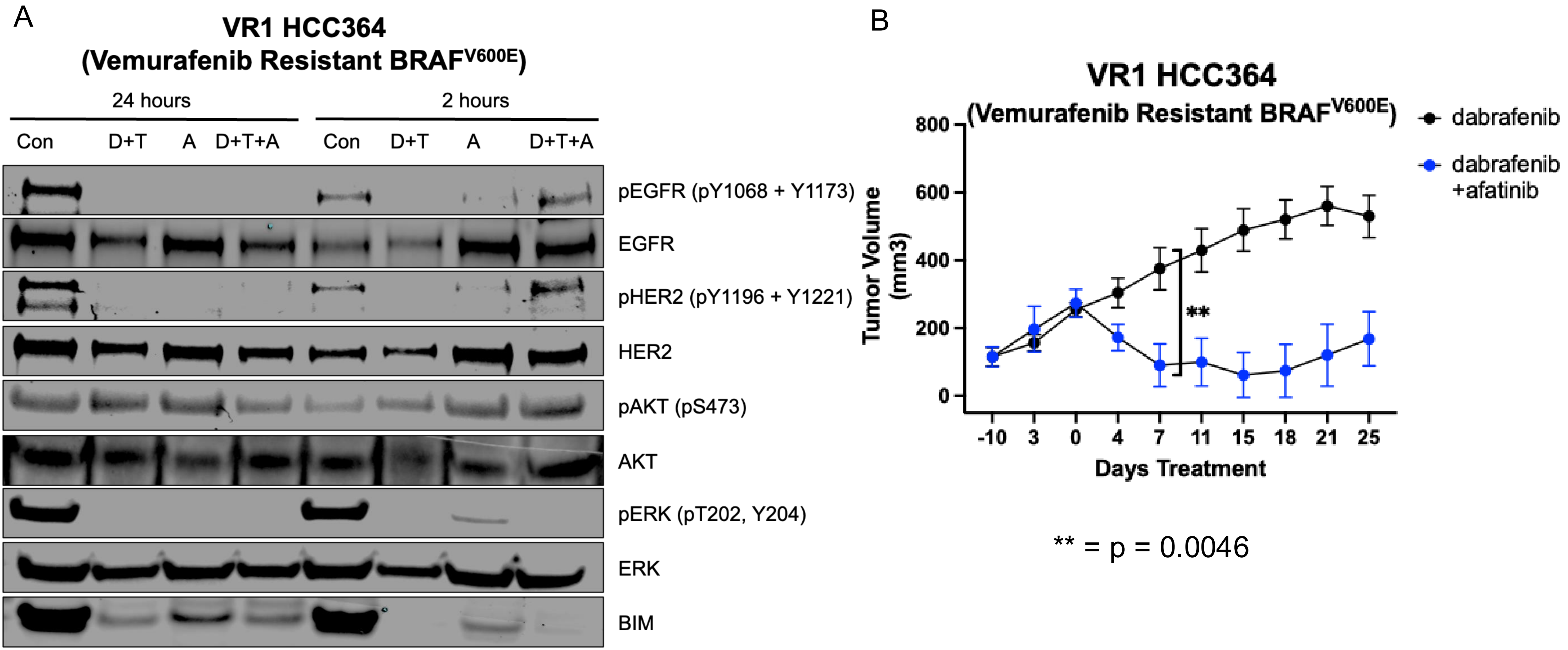


Supplementary Figure 9. Combined BRAF and pan-ERBB Pathway Targeted Therapies Are Effective in a Model of $BRAF^{V600E}$+ Lung Cancer Cell Acquired Resistance.

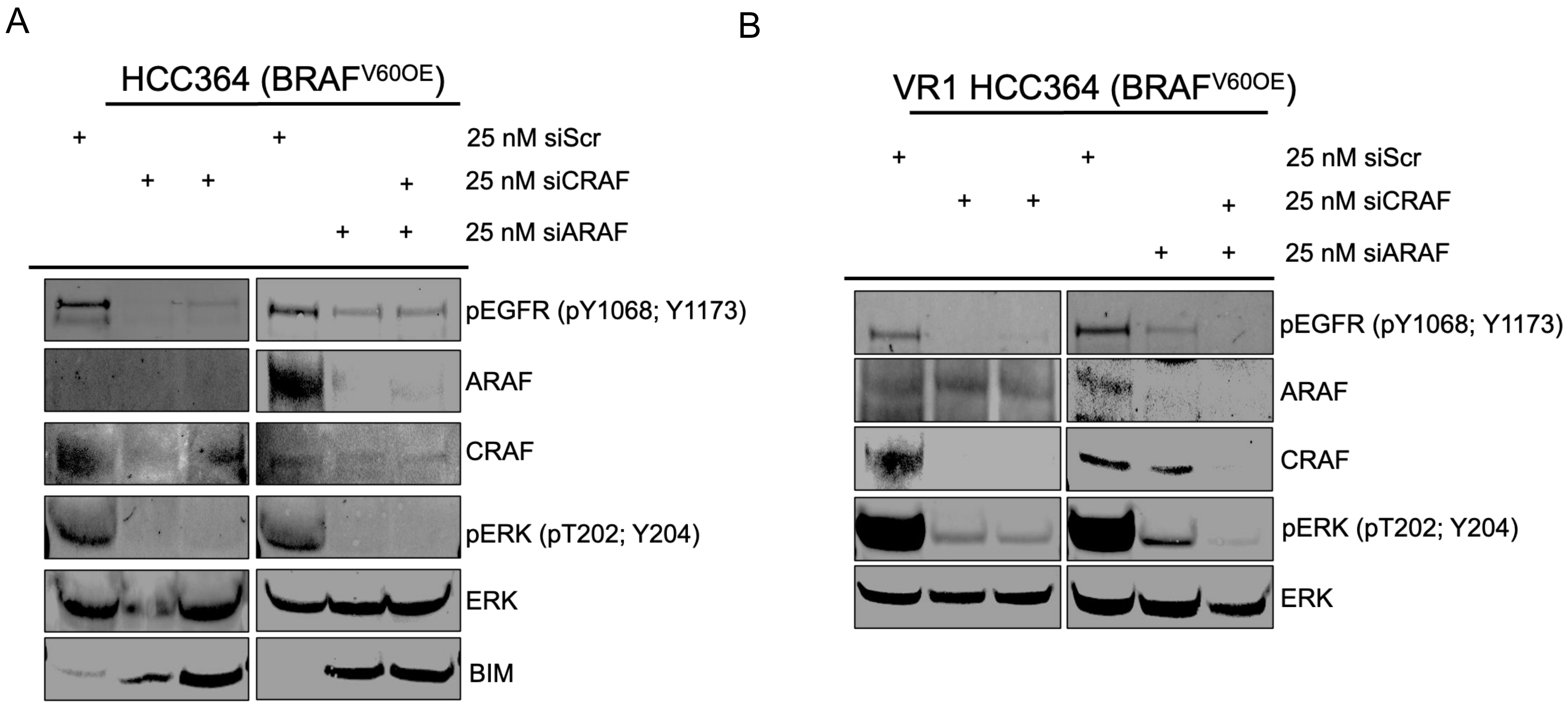


Supplementary Figure 10. RNAi Against ARAF and CRAF Phenocopy pan-ERBB Inhibition in BRAF$^{V600E}$+ Lung Cancer Models by Immunoblot Analyses.
